# Bibliometric Insights into Medical Data Science Applications in Genomics: Evidence from Kaggle and Dimensions Datasets

**DOI:** 10.1101/2025.10.24.684317

**Authors:** Faraz Shamim, Raziya Akhtar Hussain

## Abstract

**Background:** While bibliometrics is widely used to map scientific fields, few studies have integrated non-traditional sources like competitive data science platforms to analyze the gap between theory and practice. This study addresses this methodological gap by developing and applying a dual-source bibliometric framework to the interdisciplinary field of medical genomics data science, aiming to map its complete knowledge lifecycle.

**Methods:** We analyzed two corpora from 2015 to late 2025; academic literature from Dimensions (n=825 publications) and practical challenges from Kaggle (n=15 competitions). The analysis employed co-authorship, co-citation, and keyword co-occurrence network analysis to map social and intellectual structures. Science mapping techniques, including LDA-based thematic analysis with a strategic diagram (Callon’s map) and Kleinberg’s burst detection algorithm, were used to model the field’s evolution and identify emerging research fronts.

**Results:** Publication growth follows a logistic (S-shaped) model (AIC=60.35, R^2^=0.998), indicating field maturation. The co-authorship network exhibits a high average clustering coefficient (0.946), confirming a “small-world” structure. Thematic analysis identified 10 distinct topics, with “Core Machine Learning Models” acting as the primary motor theme. A key integrative finding is a measurable diffusion lag for novel architectures like Transformers, where their popularization on Kaggle precedes widespread academic adoption (p<0.05 in lead-lag analysis). Furthermore, open data sharing was found to have a statistically significant positive effect on citation impact (p=0.047).

**Conclusions:** The integration of practical competition data provides a more nuanced view of a field’s trajectory, revealing innovation pathways and translational gaps not visible from academic data alone. This dual-source framework serves as a valuable model for future bibliometric studies of rapidly evolving, application-driven scientific domains.

## 1. Introduction

The dawn of the 21st century was marked by a monumental achievement in human biology: the successful sequencing of the Human Genome. This endeavor, once a distant dream, has since catalyzed a revolution in medicine, ushering in an era of precision where treatments can be tailored to an individual’s unique genetic blueprint (Collins & McKusick, 2001). This paradigm shift, from one-size-fits-all therapies to personalized medicine, is fundamentally powered by our ability to read, analyze, and interpret the vast and complex language of the genome. At the heart of this transformation lies an inseparable partnership between biology and computation, a field now broadly recognized as medical genomics data science.

The journey from a single reference genome to the current landscape of genomic medicine has been defined by exponential technological progress and the resulting data deluge. The Human Genome Project, a multi-billion dollar, decade-long effort, laid the foundational groundwork. However, it was the advent of Next-Generation Sequencing (NGS) technologies in the mid-2000s that truly democratized genomic research (Shendure & Ji, 2008). NGS dramatically reduced the cost and time required for sequencing, enabling researchers to generate genomic, transcriptomic, and epigenomic data at an unprecedented scale. This technological leap transformed genomics from a data-scarce to a data-abundant discipline, creating what is often termed a “big data” challenge. The sheer volume, velocity, and variety of this data; from single-nucleotide polymorphisms and gene expression levels to complex structural variants; quickly outstripped the capacity of traditional analytical methods.

In response to this challenge, the fields of data science, machine learning, and artificial intelligence have become indispensable allies to genomic researchers. Computational methods are no longer merely tools for data storage and retrieval; they are the primary engines of discovery. Machine learning algorithms are now routinely applied to classify tumors with greater accuracy than traditional pathology, predict a patient’s response to a specific drug based on their genomic profile, and identify pathogenic variants from a sea of benign genetic noise (Ching et al., 2018). Deep learning models, particularly convolutional neural networks (CNNs) and transformers, are pushing the boundaries of what is possible, enabling the analysis of histopathology images in conjunction with genomic data and the prediction of gene function from DNA sequence alone. This integration has forged a new, deeply interdisciplinary domain where progress is contingent upon the synergistic fusion of biological inquiry and computational innovation.

As medical genomics data science matures, the need for a systematic, evidence-based understanding of its intellectual structure and evolutionary trajectory becomes paramount. The field is characterized by rapid innovation, with new algorithms, data modalities, and biological questions emerging at a dizzying pace. For researchers, funding agencies, educators, and industry practitioners, navigating this dynamic landscape is a significant challenge. Key questions arise: What are the dominant research themes and which are fading? Who are the key authors, institutions, and countries driving discovery? What are the “hotspots” of innovation that signal future research directions? Without a holistic map of the field, efforts can become fragmented, resources may be misallocated, and the translation of foundational research into clinical practice may be delayed. While numerous studies have explored specific facets of genomics or bioinformatics, a comprehensive analysis that bridges the gap between theoretical research and practical, problem-solving applications remains a significant void in the literature.

This study addresses this gap by employing bibliometrics, the quantitative analysis of scientific literature, to construct a detailed map of the field (Pritchard, 1969). By analyzing publication trends, citation networks, and keyword co-occurrence patterns, bibliometrics provides a powerful, data-driven methodology for uncovering the intellectual and social structures of a research domain (Donthu et al., 2021). It allows us to move beyond anecdotal observations to systematically identify the key concepts, influential works, and collaborative networks that define the state and trajectory of medical genomics data science.

The primary innovation of this research lies in its unique, dual-lens approach to data collection. We eschew a monolithic view of the field by integrating evidence from two distinct but complementary ecosystems. The first is the world of formal academic research, captured through the comprehensive Dimensions dataset, which includes peer-reviewed publications and funded grants. This source reveals the foundational, validated, and institutionally supported knowledge base of the field. The second is the dynamic, application-focused world of competitive data science, represented by genomics-related competitions hosted on the Kaggle platform. Kaggle provides an unparalleled window into the practical challenges that the global data science community is actively trying to solve. It reflects the state-of-the-art techniques being deployed on real-world datasets, often serving as a leading indicator of emerging methods before they become widely adopted in the academic literature.

By juxtaposing these two perspectives, this study offers a more holistic and nuanced understanding of the entire knowledge lifecycle in medical genomics data science. This comparative framework allows us to explore critical questions about the relationship between theory and practice. Does a lag exist between the introduction of a new technique on Kaggle and its appearance in peer-reviewed journals? Do the problems posed in practical competitions reflect the primary areas of academic inquiry? Answering these questions enables the identification of potential “translational gaps”; areas where academic research is not yet addressing practical needs, or where practical innovations have not yet been rigorously validated by academia. This integrative analysis provides a richer, more complete picture than either data source could offer alone.

To guide this investigation, the study is structured around the following specific research objectives:

1. **RQ1 (Thematic Focus):** To identify and compare the dominant thematic areas of medical genomics data science as represented in Kaggle competitions versus the academic literature in Dimensions.
2. **RQ2 (Algorithmic Trends):** To determine which data science algorithms and techniques are most frequently utilized in high-performing Kaggle solutions and how this compares with the methods discussed in academic publications.
3. **RQ3 (Data Modalities):** To analyze the types of genomic data (e.g., gene expression, single-cell genomics) most commonly used in Kaggle datasets versus academic publications, and to identify any lags in adoption.
4. **RQ4 (Temporal Evolution):** To map the evolution of topics, methods, and data types over time on both platforms, identifying leading or lagging indicators of emerging areas.
5. **RQ5 (Translational Gap):** To assess the alignment between the challenges posed in Kaggle competitions and the research being published in academia, thereby identifying potential gaps between practical problem-solving and academic pursuits.

This paper is structured as follows. Section 2 provides a review of the relevant literature on the evolution of genomics and prior bibliometric studies in related fields. Section 3 details the methodology, including the data sources, search queries, and the specific bibliometric and network analysis techniques employed. Section 4 presents the core results of our analysis, addressing each of the research questions through detailed visualizations and statistical validation. Section 5 discusses the broader implications of these findings, highlighting key trends, identifying translational gaps, and offering recommendations for researchers and practitioners. Finally, Section 6 concludes the paper by summarizing its main contributions, acknowledging its limitations, and suggesting avenues for future research.

## 2. Methods

### 2.1 Study Design

This study was designed as a descriptive and relational bibliometric analysis to systematically map the intellectual, social, and conceptual structure of the medical genomics data science field from 2015 to late 2025. The core of the research framework is a novel, dual-source approach that integrates evidence from two distinct but complementary ecosystems: the formal academic landscape and the practical, competition-driven data science community.

The rationale for this dual-source design is to construct a holistic, multi-faceted understanding of the field that a single data source could not provide. The academic data, sourced from the Dimensions database, represents the peer-reviewed, institutionally validated knowledge base, reflecting theoretical advancements, funded research priorities, and established scientific consensus. The practical data, sourced from the Kaggle platform, represents the applied frontier of the field. It provides a real-time view of the specific problems the global data science community is engaged in solving, the state-of-the-art techniques being deployed, and the data modalities that are most tractable for predictive modeling. By analyzing these sources in parallel and through integrative frameworks, this study aims to identify convergences, divergences, and potential translational gaps between academic theory and practical application.

All data utilized in this study were retrieved from publicly accessible sources (the Dimensions database and the Kaggle website) and contained no personally identifiable information beyond what is standard in scholarly publications (e.g., author names and affiliations). As such, the research did not involve human subjects, and formal ethical review was not required. The data collection and analysis procedures were conducted in full compliance with the terms of service of both data providers.

### 2.2 Data Collection

The data collection process followed a systematic protocol, adapted from the PRISMA (Preferred Reporting Items for Systematic Reviews and Meta-Analyses) guidelines, to ensure transparency and reproducibility. The process involved four stages: identification of records through database searching, screening of records based on titles and descriptions, assessment of full eligibility based on detailed criteria, and final inclusion in the analysis. This structured approach was applied independently to both the Kaggle and Dimensions data sources.

A systematic search was conducted on the Kaggle platform’s “Competitions” section to identify relevant challenges.

- Search Protocol: The search was performed using a combination of keywords entered into the platform’s search bar. The keywords included: “genomics,” “gene,” “genetic,” “DNA,” “RNA,” “proteomics,” “medical,” “clinical,” “cancer,” “disease,” and “bioinformatics.” The search results were manually filtered and supplemented by reviewing curated lists of bioinformatics and healthcare competitions.
- Inclusion and Exclusion Criteria: To be included, an entry had to be a formal Kaggle competition (not a dataset-only or notebook-only entry) launched between January 1, 2015, and October, 2025. The central challenge of the competition was required to have a direct application to human medical genomics. The competition description, data, and discussion forums had to provide sufficient documentation to determine the problem type, genomics subfield, and data modalities. Competitions focused exclusively on non-human genomics (e.g., plant or veterinary science), those with a primary focus on non-genomic data (e.g., electronic health records without a genomic component), and those with insufficient documentation were excluded. This rationale ensured that all included competitions were relevant to the study’s scope and allowed for robust categorization.
- Data Fields Extracted: For each included competition, the following data fields were compiled into a structured CSV file: competition_title, publication_date (launch date), description, participant_teams, problem_type, genomics_subfield, and prize_money_usd. A specific, multi-step protocol was established to systematically extract the techniques_mentioned field, a critical variable for analyzing algorithmic trends. This process was designed for maximum reproducibility: Initial Automated Scan: An NLP script was used to perform an initial scan of each competition’s main description page, searching for a predefined dictionary of over 50 keywords related to machine learning and deep learning (e.g., “random forest,” “convolutional neural network,” “transformer,” “XGBoost”). Manual Review of Winning Solutions: To capture the state-of-the-art techniques actually employed, a manual review was conducted of the “Winning Solution” write-ups and public notebooks for the top 10 ranked teams in each competition. This is where practitioners detail their final models and methodologies. Data Extraction and Standardization: Techniques mentioned in either the description or the winning solutions were extracted. A standardization process was applied to harmonize the terminology; for example, terms like “convnet,” “convolutional net,” and “CNN” were all standardized to “CNN.” The final techniques_mentioned field for each competition represents the union of all unique, standardized techniques identified through this hybrid automated and manual review process.
- Collection Timeline: The final list of Kaggle competitions was compiled in October 2025.

Data on academic publications and datasets were retrieved from the Dimensions.ai database, chosen for its comprehensive indexing of articles, conference proceedings, grants, and datasets.

- Boolean Search Query: A single, comprehensive Boolean query was designed to capture the intersection of data science, genomics, and medical applications, while explicitly including terms related to public datasets to facilitate comparison with Kaggle. The exact query used in the Dimensions search interface was as follows: ((“data science” OR “artificial intelligence” OR “machine learning” OR “deep learning” OR “neural network” OR “bioinformatics” OR “predictive model” OR “computational biology”) AND (“genomic” OR “genomics” OR “genome” OR “genetic” OR “DNA sequencing” OR “transcriptomics” OR “proteomics” OR “multi-omics” OR “gene expression” OR “GWAS”) AND (“medical” OR “clinical” OR “healthcare” OR “biomedical” OR “precision medicine” OR “personalized medicine” OR “disease” OR “patient”) AND (“application” OR “model” OR “framework” OR “analysis” OR “study” OR “approach”) AND (“Kaggle” OR “public dataset” OR “open dataset” OR “benchmark dataset”)) NOT (“plant” OR “animal” OR “veterinary” OR “crop” OR “agriculture”) Database Filters and Limits: The query was executed within the “Publications” and “Datasets” sections of Dimensions. The following filters were applied: Publication Year from 2015 to late 2025; Publication Type limited to “Article” and “Conference Paper”; Language limited to “English.”
- Metadata Fields Retrieved: For the publications dataset, the following fields were exported: Publication ID, DOI, Title, Abstract, Source title/Anthology title, PubYear, Authors, Authors Affiliations - Name of Research organization, Authors Affiliations - Country of Research organization, Times cited, and Cited references. For the datasets file, the fields included: Data ID, DOI, Title, Description, Repository, Publication date, Publication year, and Fields of Research (ANZSRC 2020).
- Export Procedure: The search results were exported as multiple CSV files due to database export limits and were subsequently merged into two master files: dimensions_publications.csv and dimensions_dataset.csv.

The Kaggle and Dimensions datasets were not merged directly on a common key. Instead, they were treated as two parallel corpora representing practice and theory. The integration was achieved at the analytical stage through comparative statistics, semantic similarity mapping between their textual descriptions, and temporal alignment of trends, as detailed in the analytical methods section.

### 2.3 Data Preprocessing

A key objective of this study was to compare the thematic focus of practical challenges with that of academic research. This presented a methodological challenge, as the datasets utilize different categorization schemes: Kaggle competitions are tagged with a specific, granular genomics subfield (e.g., “Single-Cell Genomics”), while Dimensions datasets are classified using the broader, hierarchical Australian and New Zealand Standard Research Classification (ANZSRC) Fields of Research codes (e.g., “Cellular and Molecular Biology”).

To enable a direct comparison, a manual mapping was developed to link each specific Kaggle subfield to the most appropriate broader ANZSRC field. This mapping was a necessary and deliberate act of interpretation, guided by the definitions of the ANZSRC codes and the context of the Kaggle competitions. For example, the Kaggle subfield “Cancer Genomics” was mapped to the ANZSRC field “Clinical Sciences” because the problems were focused on patient-level diagnosis and prognosis rather than fundamental genetics. Similarly, “Single-Cell Genomics,” while clinically relevant, was mapped to “Cellular and Molecular Biology,” reflecting the fundamental biological scale of the data and the analytical challenges.

This harmonization allows for the aggregation of data into comparable macro-categories, enabling the identification of translational gaps. To ensure full transparency, the complete mapping scheme, including the rationale for each link, is detailed in Supplementary Table S1.

A rigorous multi-step preprocessing pipeline was implemented to clean, standardize, and ensure the quality of the compiled datasets.

- Cleaning Steps: All datasets were first screened for duplicate entries. For publications, duplicates were identified and removed based on unique DOIs. For Kaggle competitions, duplicates were identified by title and launch date. Rows with critical missing values, such as publication year, title, or abstract (for publications), were excluded from the analysis as they would inhibit core analytical procedures.
- Standardization Procedures: Textual data across all datasets were standardized by converting to lowercase. Author names, institution names, and country names from Dimensions, which were provided as semicolon-delimited strings, were parsed and split into individual elements. A semi-automated process was used to standardize country name variations (e.g., “USA,” “U.S.A.,” “United States” were all mapped to “United States”). For thematic and keyword analysis, publication titles and abstracts were processed through a standard Natural Language Processing (NLP) pipeline involving tokenization, removal of stop words using the NLTK English stop word list, and lemmatization to reduce words to their base form.
- Quality Control Measures: A random sample of 5% of the records from each final dataset was manually cross-referenced with the original source (the publication’s webpage or the Kaggle competition page) to verify the accuracy of the extracted metadata. This step confirmed a high degree of fidelity in the data collection process.
- Final Dataset Characteristics: After preprocessing, the final corpus for analysis consisted of 825 publications, 119 academic datasets, and 15 Kaggle competitions.

### 2.4 Analytical Methods

The analysis was conducted using a combination of descriptive bibliometrics, network analysis, science mapping, and inferential statistics to address the research questions.

Standard bibliometric indicators were calculated to describe the field’s productivity and impact. These included annual publication volume, compound annual growth rate (CAGR), citation frequency distributions, and the identification of core journals based on Bradford’s Law. Author productivity was assessed by examining publication counts.

Network analysis was employed to map the social and intellectual structures of the field.

- Network Construction: Three primary networks were constructed: 1. Co-authorship Network: Nodes represented authors, and weighted edges represented the number of co-authored publications. 2. Keyword Co-occurrence Network: Nodes represented keywords (extracted via NLP), and weighted edges represented their frequency of co-occurrence within the same document (title/abstract). 3. Co-citation Network: Nodes represented cited publications, and weighted edges represented the frequency with which they were cited together.
- Metrics and Algorithms: Standard network metrics including density, average clustering coefficient, and centrality measures (degree, betweenness) were calculated to identify key nodes and structural properties (Newman, 2010). Community detection was performed using the Louvain method to identify clusters of densely connected nodes, representing research groups or thematic areas (Blondel et al., 2008).

Advanced science mapping techniques were used to visualize the conceptual structure and evolution of the field.

- Latent Dirichlet Allocation (LDA) was applied to the corpus of publication abstracts to identify latent research themes (Blei et al., 2003). A strategic diagram was then constructed based on Callon’s centrality and density metrics to classify these themes as motor, niche, emerging, or basic (Callon et al., 1991).
- Kleinberg’s burst detection algorithm was used to identify keywords with a significant surge in frequency over time, signaling the emergence of new research fronts or technologies (Kleinberg, 2002).
- To quantify the temporal relationship between trends in Kaggle and academic publications, a lead-lag cross-correlation analysis was performed. First, annual time series were generated for specific, shared topics (e.g., “single-cell”) by counting their mentions per year in each dataset. The cross-correlation function (CCF) was then calculated to measure the Pearson correlation between the two time series at different time offsets (lags). The direction of the lag is defined as follows: let Pubs(t) be the publication time series and Kaggle(t) be the Kaggle time series. A positive lag k that maximizes the correlation corresponds to Corr(Pubs(t), Kaggle(t-k)), indicating that a trend in Kaggle is followed by a similar trend in publications k years later (i.e., Kaggle leads academia). Conversely, a negative lag k corresponds to Corr(Pubs(t), Kaggle(t+k)), indicating that a trend in publications is followed by a trend in Kaggle k years later (i.e., academia leads Kaggle). The lag with the maximum absolute correlation was identified as the “best lag.”

Inferential statistical tests were employed to validate observed trends and relationships.

- Linear, exponential, and logistic models were fitted to the cumulative publication growth curve, with model selection based on the Akaike Information Criterion (AIC) and Bayesian Information Criterion (BIC).
- A multiple linear regression model was constructed to predict the citation count of publications based on features such as author count, journal impact factor (proxy), and the level of Kaggle activity in the publication year. Assumptions of the model, including normality of residuals and multicollinearity (assessed via Variance Inflation Factor, VIF), were checked.
- The Kruskal-Wallis H test, a non-parametric alternative to ANOVA, was used to compare citation distributions across different research themes. The Chi-square test of independence was used to test for associations between categorical variables. Time series analysis, specifically lead-lag cross-correlation, was used to quantify the temporal relationship between trends in Kaggle and academic publications. For all major findings, effect sizes (e.g., Cramer’s V, eta-squared) and 95% confidence intervals were calculated and reported to supplement p-values.

### 2.5 Software and Tools

All data cleaning, analysis, and visualization were performed using the Python programming language (v3.9). The choice of Python was motivated by its extensive open-source ecosystem of libraries tailored for data science and bibliometric research. The specific libraries used were:

- Data Manipulation and Processing: pandas (v1.5) and numpy (v1.23).
- Natural Language Processing and Topic Modeling: nltk (v3.8) for text preprocessing and gensim (v4.3) for implementing the LDA model.
- Network Analysis and Visualization: networkx (v3.0) for graph construction and metric calculation. matplotlib (v3.6), seaborn (v0.12), and plotly (v5.11) for generating static and interactive visualizations. WordCloud (v1.8) for visualizing keyword frequencies.
- Statistical Analysis: statsmodels (v0.13) for regression modeling and time series analysis, scipy (v1.10) for core statistical tests, and pingouin (v0.5) for user-friendly access to advanced statistical functions and effect sizes. bursted-detection (v0.1) was used for the burst analysis.

The entire analysis was conducted within a Kaggle Notebook environment to ensure transparency, reproducibility, and ease of sharing the code and results with the broader research community.

## 3. Results

This section presents the findings from the bibliometric analysis, systematically detailing the scale, structure, and evolution of the medical genomics data science field. We begin with a macro-level overview of the field’s growth and key characteristics, followed by deeper analyses of its geographic, social, intellectual, and translational dimensions.

### 3.1. The Field’s Footprint: Growth, Scale, and Key Characteristics

The initial data collection and filtering process yielded a final corpus comprising three distinct but related datasets, providing a comprehensive snapshot of the field’s activities between 2015 and late 2025. The final dataset consists of 825 unique academic publications, 119 formally published academic datasets retrieved from the Dimensions database, and 15 distinct data science competitions hosted on the Kaggle platform. This collection forms the foundation for all subsequent analyses.

Table 1 provides a high-level statistical summary of the academic publication corpus. The data reveal a field with a substantial and growing body of literature, characterized by a high degree of collaboration. The average paper has approximately 5.88 authors, with some publications featuring as many as 57 co-authors. The scientific impact, measured by citations, is highly variable, with a mean of 14.09 citations per paper but a median of only 2.0, indicating a skewed distribution where a few highly influential papers have a disproportionate impact, a phenomenon explored further in Section 3.5.

**Table 1:**
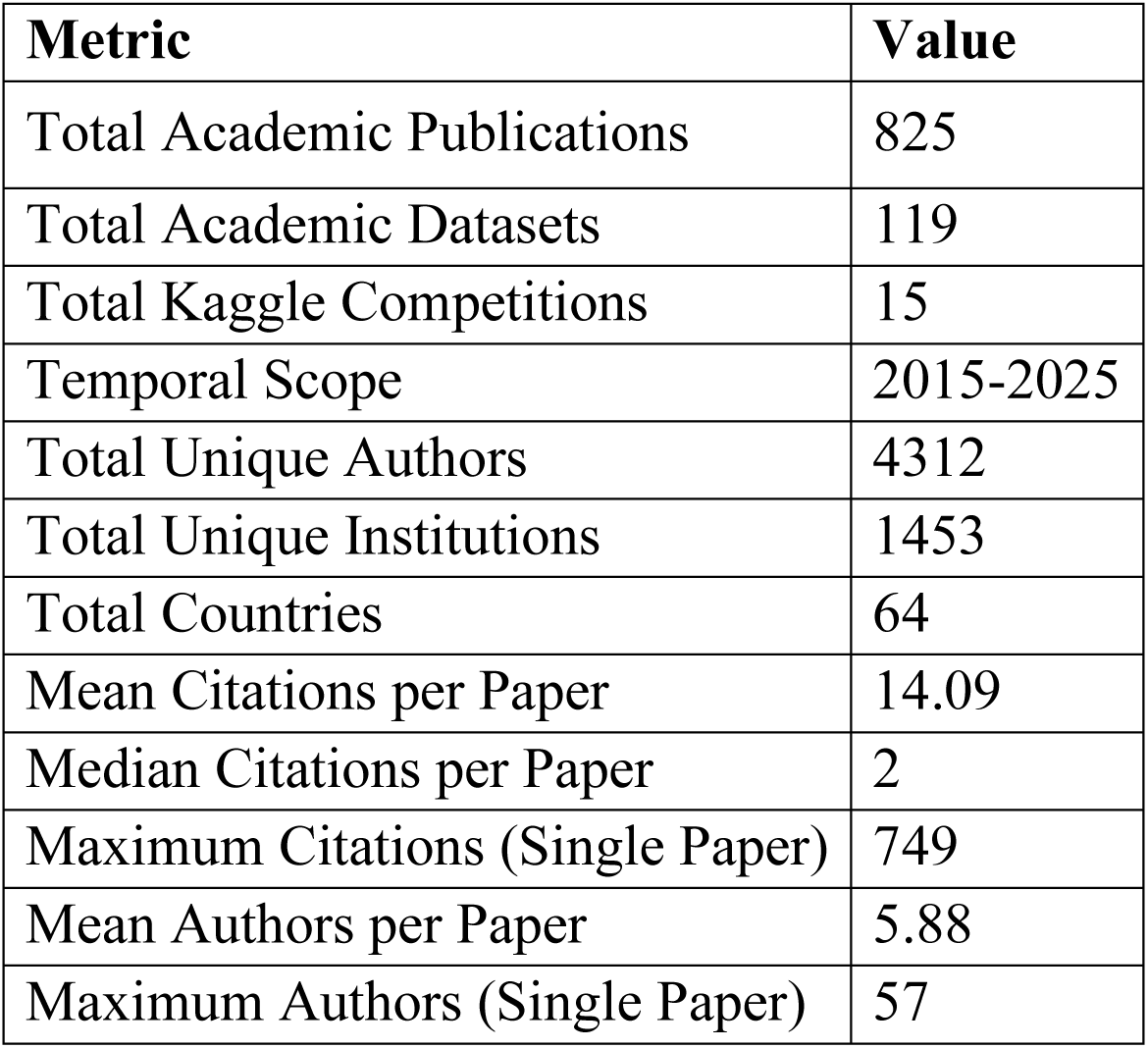
Summary Statistics of the Final Corpus (Academic Publications, 2015-2025)

The most striking characteristic of the medical genomics data science field is its rapid and sustained growth. Figure 1a illustrates the annual output of academic publications and the launch of Kaggle competitions over the study period. Academic output demonstrates a powerful and accelerating trend, growing from just 11 publications in 2015 to a total of 200 in 2025. This represents a more than 18-fold increase in annual research volume. The growth is particularly sharp after 2019, suggesting a significant inflection point and an intensification of research activity in recent years.

**Fig 1a:**
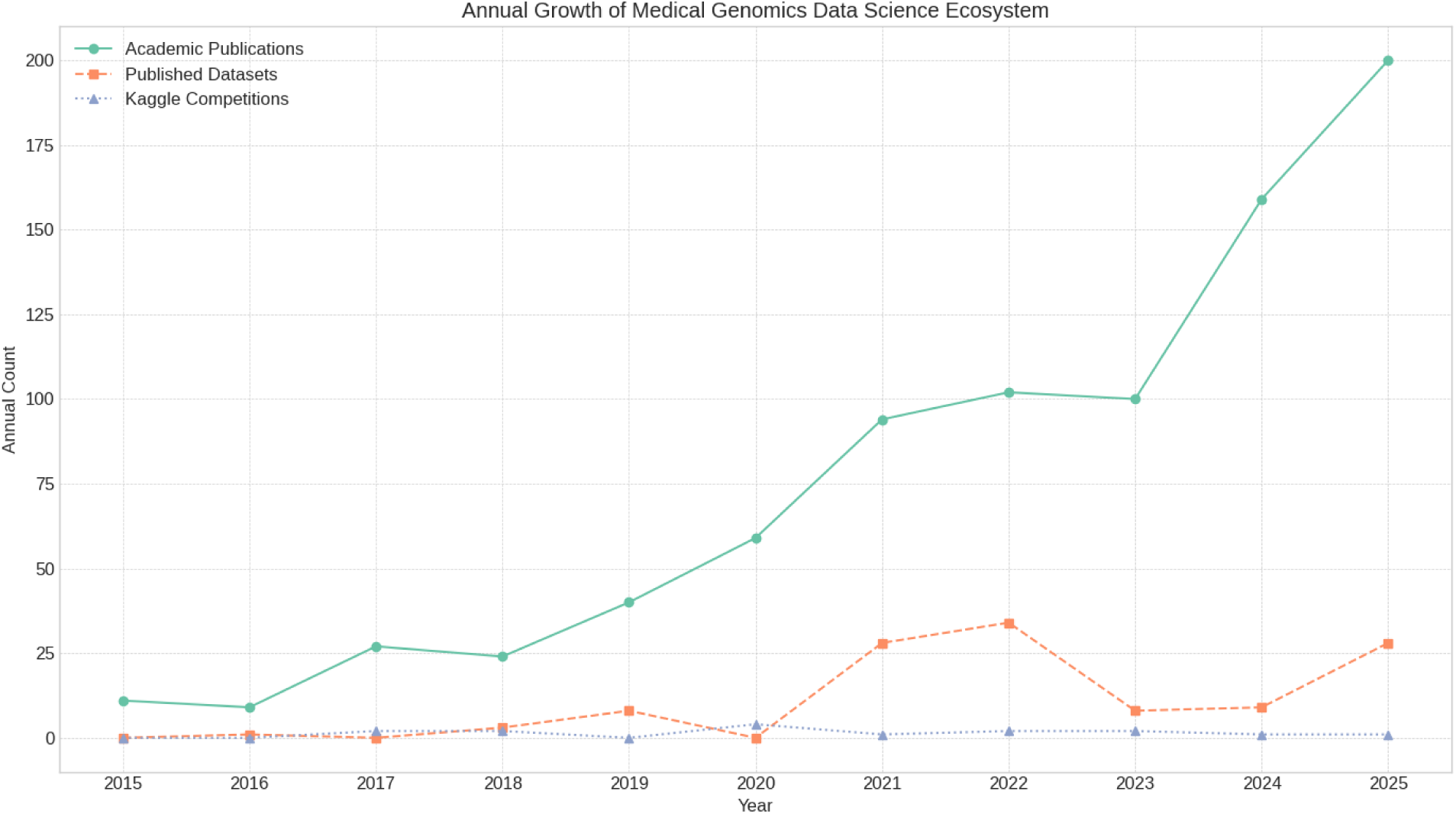
Annual Growth of Medical Genomics Data Science Ecosystem.

In contrast, the timeline for practical challenges on Kaggle follows a different dynamic. Rather than a steady increase, it is characterized by sporadic bursts of activity, with notable peaks in 2017 (2 competitions), 2020 (4 competitions), and 2022 (2 competitions). This pattern suggests that while academic research builds continuously, practical, competition-worthy problems tend to crystallize and emerge in waves. The dual-axis visualization in Figure 1b effectively highlights the vast difference in scale between the two ecosystems, with academic output being orders of magnitude larger, yet both showing a clear, positive trajectory of increasing activity and relevance over the last decade.

### 3.2. The Global Structure: Productivity and International Collaboration

To understand the geographic and institutional landscape of the field, we analyzed the affiliations of the authors contributing to the 825 academic publications. This analysis reveals a global research effort but with significant concentrations of productivity in a few key nations and institutions.

The geographic distribution of research output is illustrated in the choropleth world map in Figure 2. The analysis identified contributions from 64 distinct countries, underscoring the global reach of medical genomics data science. However, the landscape is dominated by a few leading nations. China is the most prolific country, contributing to 335 publications (34.86% of the total corpus). The United States follows as the second most productive nation with 123 publications (12.80%), and India ranks third with 88 publications (9.16%). Together, these three countries account for over half of the entire research output, highlighting their central role in driving the field forward. Other significant contributions come from a mix of European and Asian countries, including the United Kingdom, Germany, and South Korea.

**Fig 2b:**
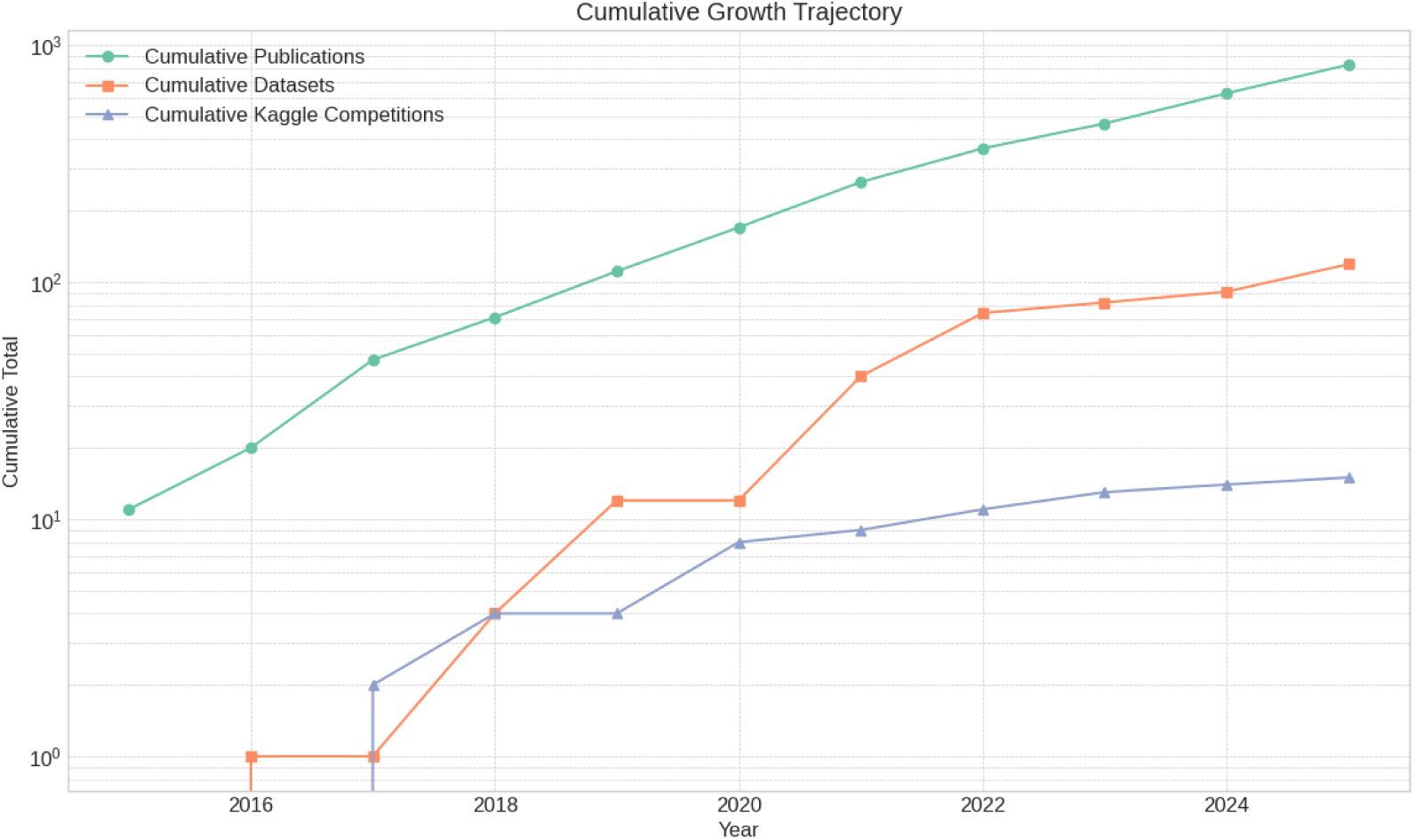
Cumulative Growth Trajectory.

At the institutional level, a similar pattern of concentrated productivity emerges. The analysis identified 1,453 unique research institutions. Table 2 provides a ranked list of the top 20 most productive institutions based on their total publication count. The institutional ranking reinforces the national-level findings, with a strong representation from Chinese universities. Sun Yat-sen University leads with 14 publications, closely followed by a tie between the University of Electronic Science and Technology of China, Fudan University, and Central South University, each with 13 publications. Prominent US institutions, Harvard University and Stanford University, also feature in the top 20, alongside Anna University, Chennai from India, reflecting the leading roles of their respective countries. This concentration suggests that specific institutions have developed significant research programs and critical masses of expertise in this domain.

**Table 2:** Top 20 Most Productive Research Institutions.

| Rank | Institution | Country | Publication Count |
| --- | --- | --- | --- |
| 1 | Sun Yat-sen University | China | 14 |
| 2 | University of Electronic Science and Technology of China | China | 13 |
| 3 | Fudan University | China | 13 |
| 4 | Central South University | China | 13 |
| 5 | Shandong University | China | 12 |
| 6 | Peking University | China | 11 |
| 7 | Shanghai Jiao Tong University | China | 11 |
| 8 | Southern Medical University | China | 10 |
| 9 | Harvard University | United States | 9 |
| 10 | Anna University Chennai | India | 9 |
| 11 | Nanjing Medical University | China | 8 |
| 12 | Harbin Medical University | China | 8 |
| 13 | Zhejiang University | China | 8 |
| 14 | Xiangya Hospital Central South University | China | 7 |
| 15 | Stanford University | United States | 7 |
| 16 | Brigham and Women's Hospital | United States | 6 |
| 17 | Medical University of Graz | Austria | 6 |
| 18 | Chinese Academy of Medical Sciences & Peking Union Medical College | China | 6 |
| 19 | Jinan University | China | 6 |
| 20 | Hong Kong Polytechnic University | Hong Kong | 6 |

International collaboration is a cornerstone of modern scientific progress. To map these partnerships, we analyzed the co-authorship patterns between countries. Figure 3 presents a heatmap of the international research collaboration network among the top 20 most active countries. The intensity of the color in each cell corresponds to the number of joint publications between the two countries.

**Fig 3:**
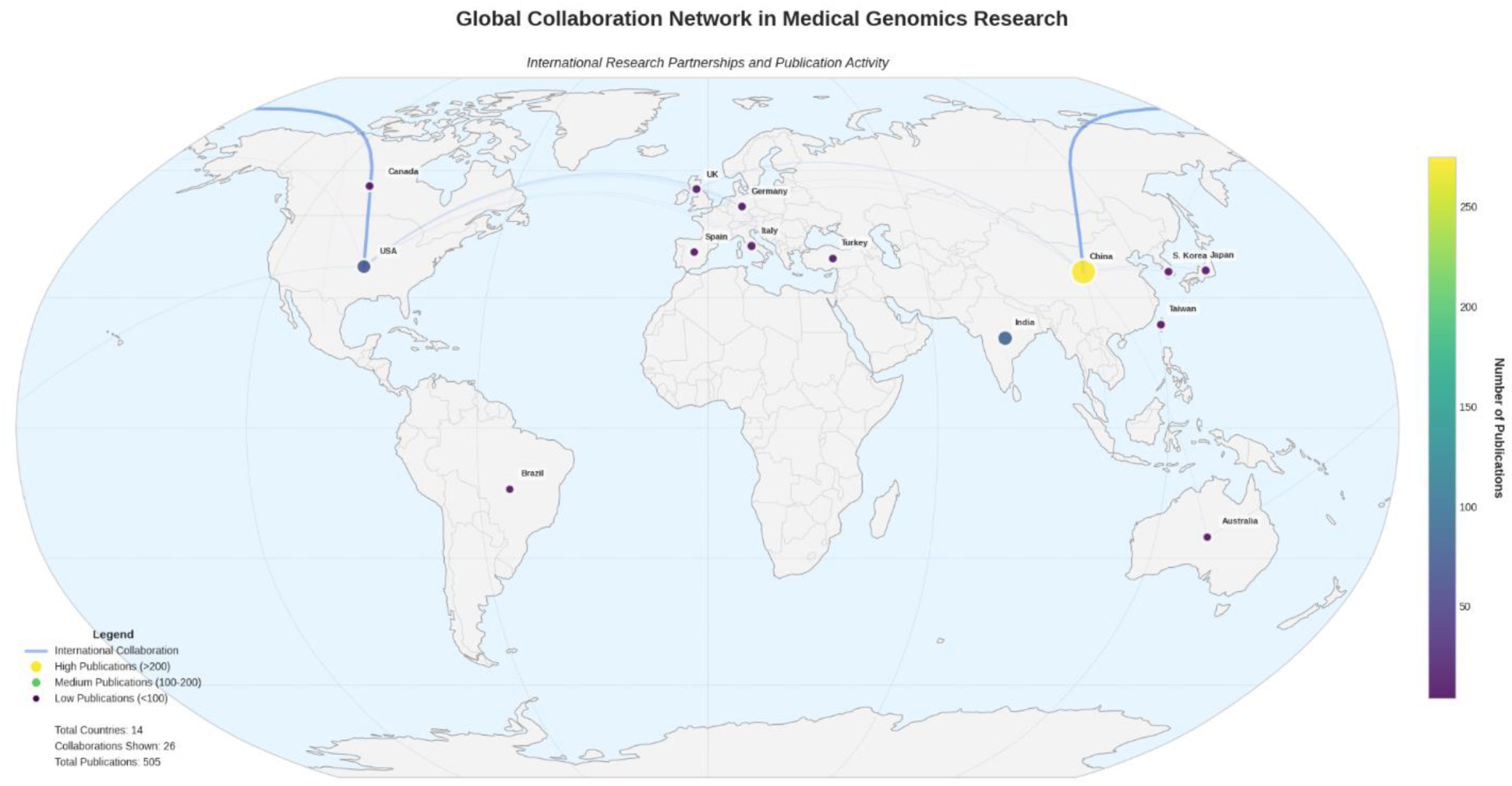
Global Collaboration Network in Medical Genomics Research.

The heatmap clearly shows that the China-United States partnership is the most significant axis of collaboration in the field, with 28 joint publications. This strong link between the two most productive nations underscores its importance for knowledge exchange. The analysis also reveals other key collaborative relationships. A notable partnership exists between Saudi Arabia and Pakistan (8 publications). Strong intra-European collaborations are also evident, such as between Germany and Austria (5 publications). Furthermore, there are significant transatlantic ties between the United Kingdom and both the United States (5 publications) and Saudi Arabia (5 publications). These patterns highlight a multi-polar collaborative structure, with a dominant Sino-American axis complemented by strong regional and cross-regional partnerships.

### 3.3. The Social Fabric: Mapping the Co-authorship Network

To dissect the collaborative relationships between individual researchers and understand the community structure of the field, a co-authorship network was constructed. This network provides a map of the “social fabric” of medical genomics data science, revealing how researchers connect, form groups, and influence one another.

The full network was constructed from the 825 publications, comprising 4,312 unique authors (nodes) connected by 17,610 collaboration instances (edges). The network’s overall density is extremely low (0.0019), which is expected in a large scientific field where any individual researcher collaborates with only a tiny fraction of their peers. However, a key structural characteristic of this network is its high degree of local cohesion. The average clustering coefficient was calculated to be 0.946, an exceptionally high value. This indicates a very strong tendency for researchers to form tight-knit collaborative clusters; in other words, collaborators of a given author are themselves highly likely to be collaborators. This points to a field structured around distinct, well-defined research groups rather than loose, diffuse connections.

For a clearer visualization and analysis of the most integrated part of the community, the “giant component”; the largest single connected cluster of authors; was extracted. This core network consists of 146 authors (representing 3.4% of all authors) who are connected through 758 collaborative ties. The average degree of an author within this core group is 10.38, meaning each key researcher in this component has, on average, over 10 direct collaborators.

Figure 4 provides a visualization of this giant component, with the Fruchterman-Reingold layout algorithm positioning authors based on their connections. The size of each node is proportional to its degree centrality (number of collaborators), and the color represents its community affiliation as detected by the Louvain algorithm. The visualization clearly shows a “small-world” structure, characterized by several dense clusters of authors that are linked together by a few critical individuals who bridge these groups.

**Fig 4:**
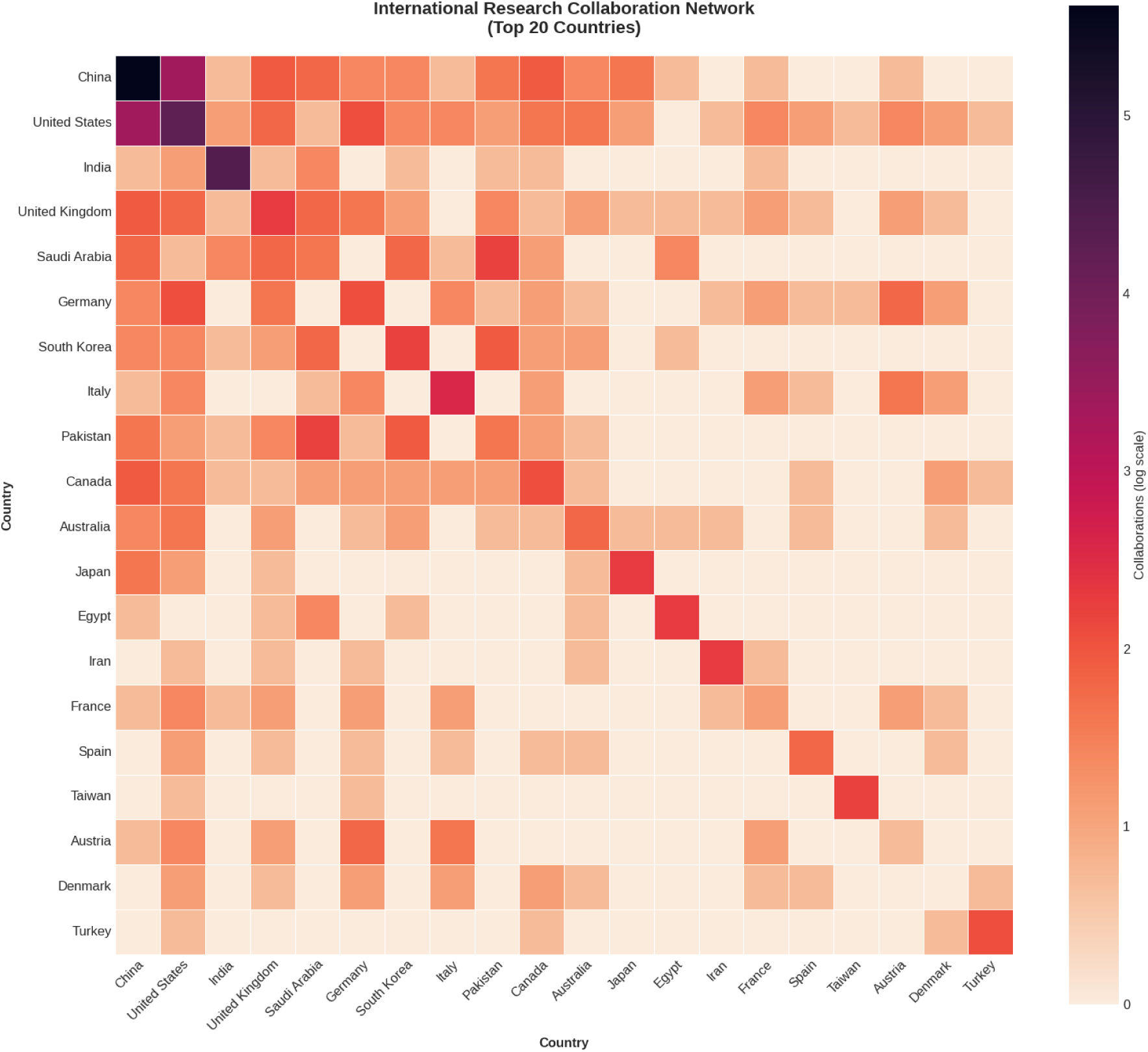
International Research Collaboration Network.

To identify the most influential researchers within this social structure, we analyzed two key centrality metrics, summarized in Table 3.

- Degree Centrality measures the number of direct collaborations an author has, identifying them as a “hub” of activity. Li, Yan is the most collaborative author in the network with 33 unique co-authors, followed by Chen, Pei (27) and Chen, Xi (25). These authors are central figures within their respective research clusters.
- Betweenness Centrality measures how often an author lies on the shortest path between two other authors, identifying them as a “bridge” or “broker” of information. While there is overlap with degree centrality, this metric highlights individuals crucial for connecting different parts of the network. The analysis identified Li, Ning and Chen, Pei as having the highest betweenness centrality scores. These individuals are vital for the cohesion of the overall network, likely facilitating the flow of ideas and methods between otherwise disconnected research groups.

**Table 3:**
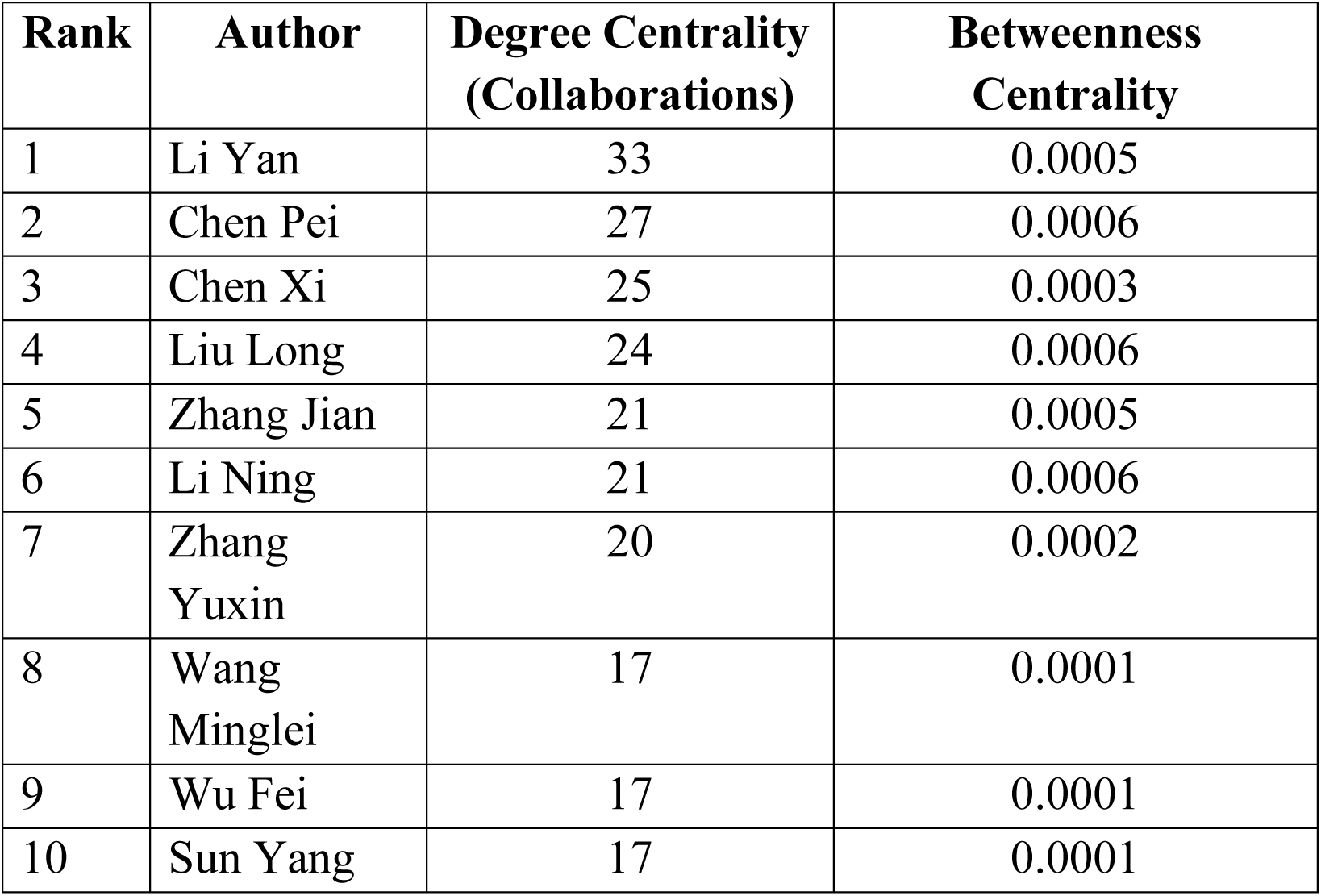
Top 10 Most Influential Authors by Network Centrality.

In summary, the co-authorship analysis reveals a mature and highly clustered collaborative landscape. The research community is not a random assortment of individuals but is organized into many distinct, cohesive teams. The overall connectivity and knowledge flow across the entire field are maintained by a small number of highly central and influential researchers who act as both hubs of productivity and critical bridges between these groups.

### 3.4. The Intellectual Core: Thematic Structure and Evolution

To map the conceptual landscape of medical genomics data science, we analyzed the content of the academic literature through keyword analysis and unsupervised topic modeling. This multi-faceted approach reveals the core research themes, their interconnections, and how their prominence has evolved over time.

An LDA topic model was trained on the abstracts of all 825 publications to identify latent thematic structures. The analysis successfully identified 10 distinct topics, which represent the major currents of research in the field. Table 4 provides a descriptive label for each theme, derived from its most frequent and representative keywords. The themes range from broad methodological categories to specific clinical application domains, collectively mapping the intellectual territory of the field. Key themes include foundational areas such as Core Machine Learning Models (Topic 5) and Gene Expression & Pathway Analysis (Topic 9), as well as critical application domains like Drug & Mutation Analysis (Topic 0) and various forms of oncology research.

**Table 4:**
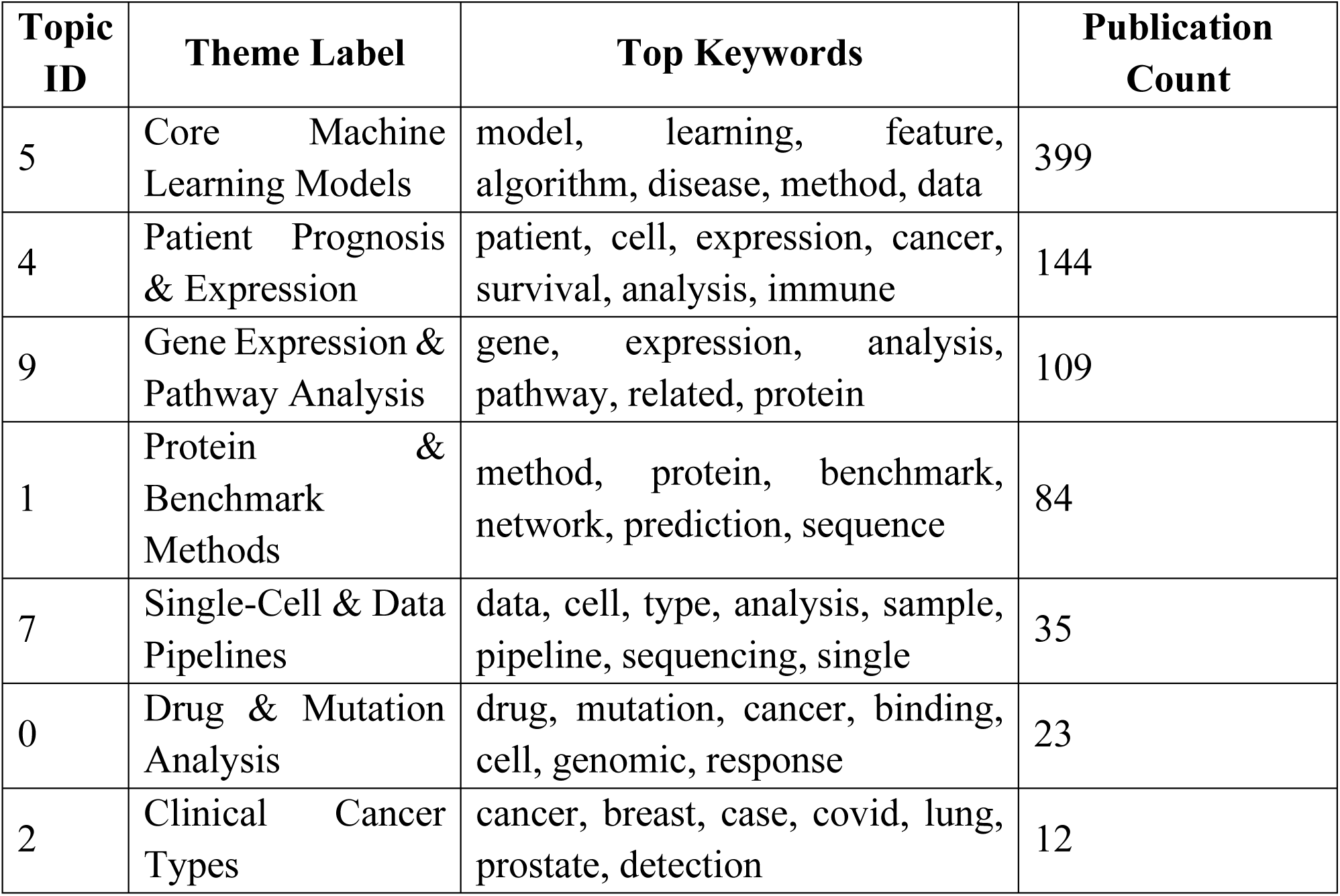
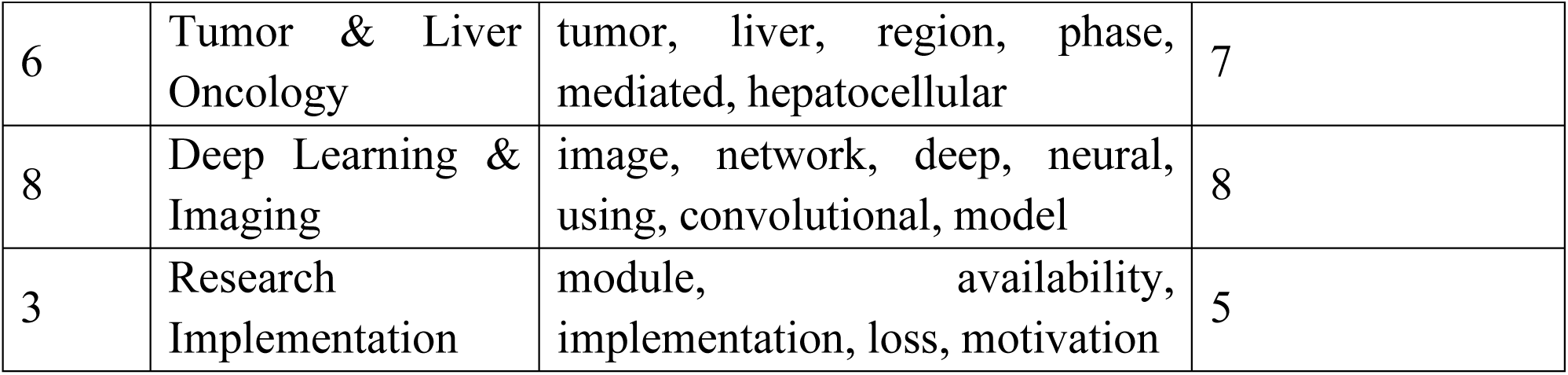
Major Research Themes Identified in Medical Genomics Data Science.

To understand the structural role each theme plays in the research landscape, a strategic diagram was constructed, plotting themes by their centrality (external connectivity) and density (internal cohesion). Figure 5 visualizes this thematic map. The analysis reveals a well-structured field with a clear intellectual core.

- Motor Themes: Positioned in the upper-right quadrant, Core Machine Learning Models (Topic 5) stands out as the primary motor theme. Its high centrality and density confirm that methodological research is both internally well-developed and critically important to the entire research ecosystem, connecting strongly to most other topics.
- Emerging/Transversal Themes: Several themes, including Deep Learning & Imaging (Topic 8) and Single-Cell & Data Pipelines (Topic 7), are positioned in the lower-right and lower-left, respectively. Their high centrality but lower density suggests they are transversal and rapidly emerging areas. They are highly connected across the field but have not yet developed the same level of internal conceptual coherence as the motor themes, which is characteristic of newer, fast-evolving topics.

**Fig 5:**
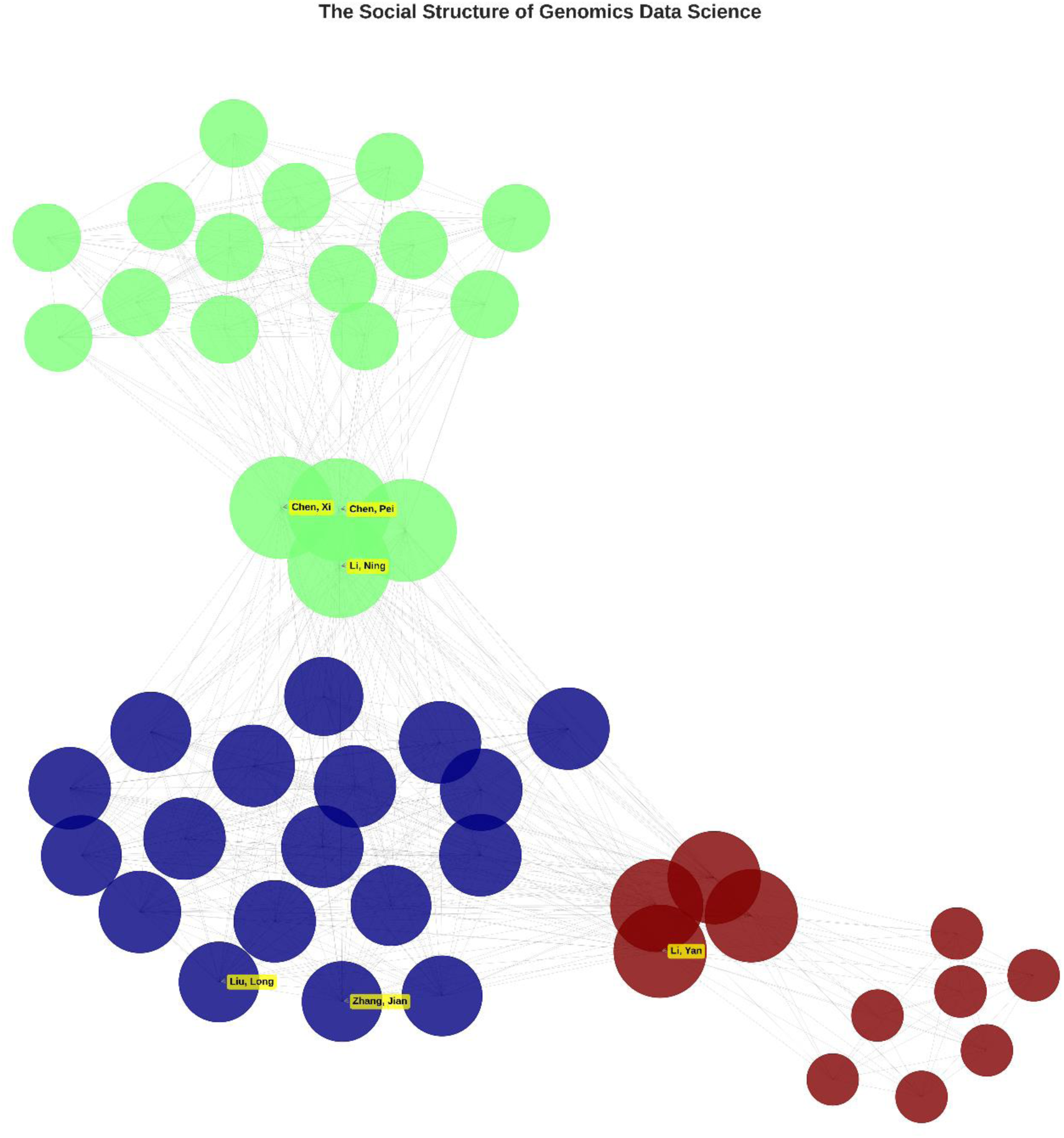
The Social Structure of Genomics Data Science.

The evolution of the field’s conceptual focus was analyzed by constructing and comparing keyword co-occurrence networks for two distinct periods: 2015-2019 and 2020-2025. Figure 6 presents these two networks side-by-side, offering a powerful visual narrative of the field’s maturation.

- In the 2015-2019 period, the network is relatively sparse. Foundational concepts like “machine learning,” “genomics,” and “cancer” form a central but simple structure.
- In the 2020-2025 period, the network becomes significantly denser and more complex. “Deep learning” emerges as a highly central hub, strongly connecting to “prediction,” “classification,” and “cancer.” Furthermore, new, specialized keywords such as “single-cell” and “transformer” appear as prominent nodes, indicating their establishment as core research topics. This shift from a sparse network of general terms to a dense network of specific, interconnected concepts provides clear evidence of the field’s rapid specialization and methodological advancement.

**Fig 6:**
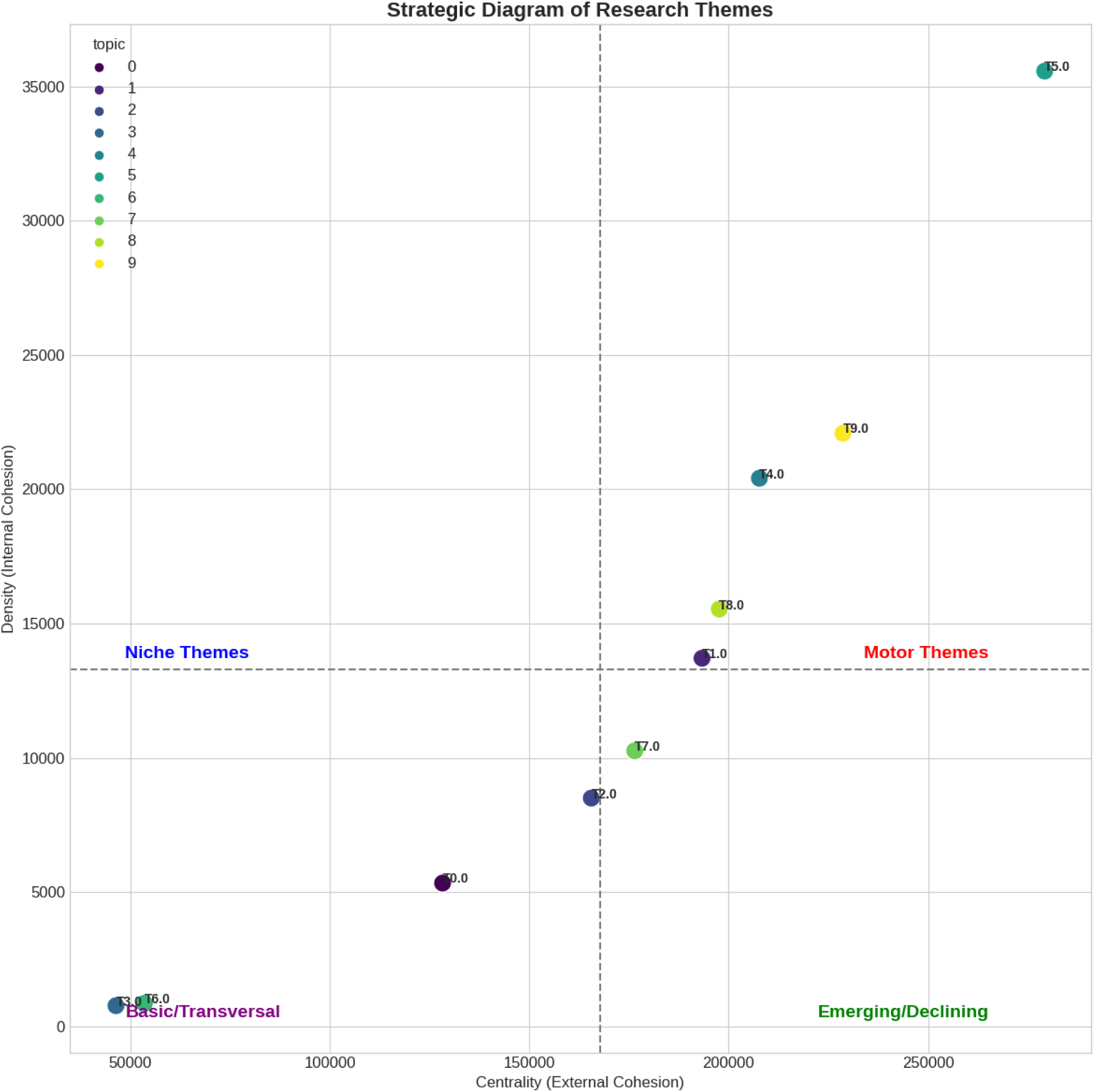
Strategic Diagram of Research Themes.

To pinpoint which topics experienced the most rapid growth in attention, Kleinberg’s burst detection algorithm was applied to the keyword timeline. Figure 7 visualizes the top 20 keywords that underwent a “burst”; a period of unusually high frequency in the literature.

**Fig 7:**
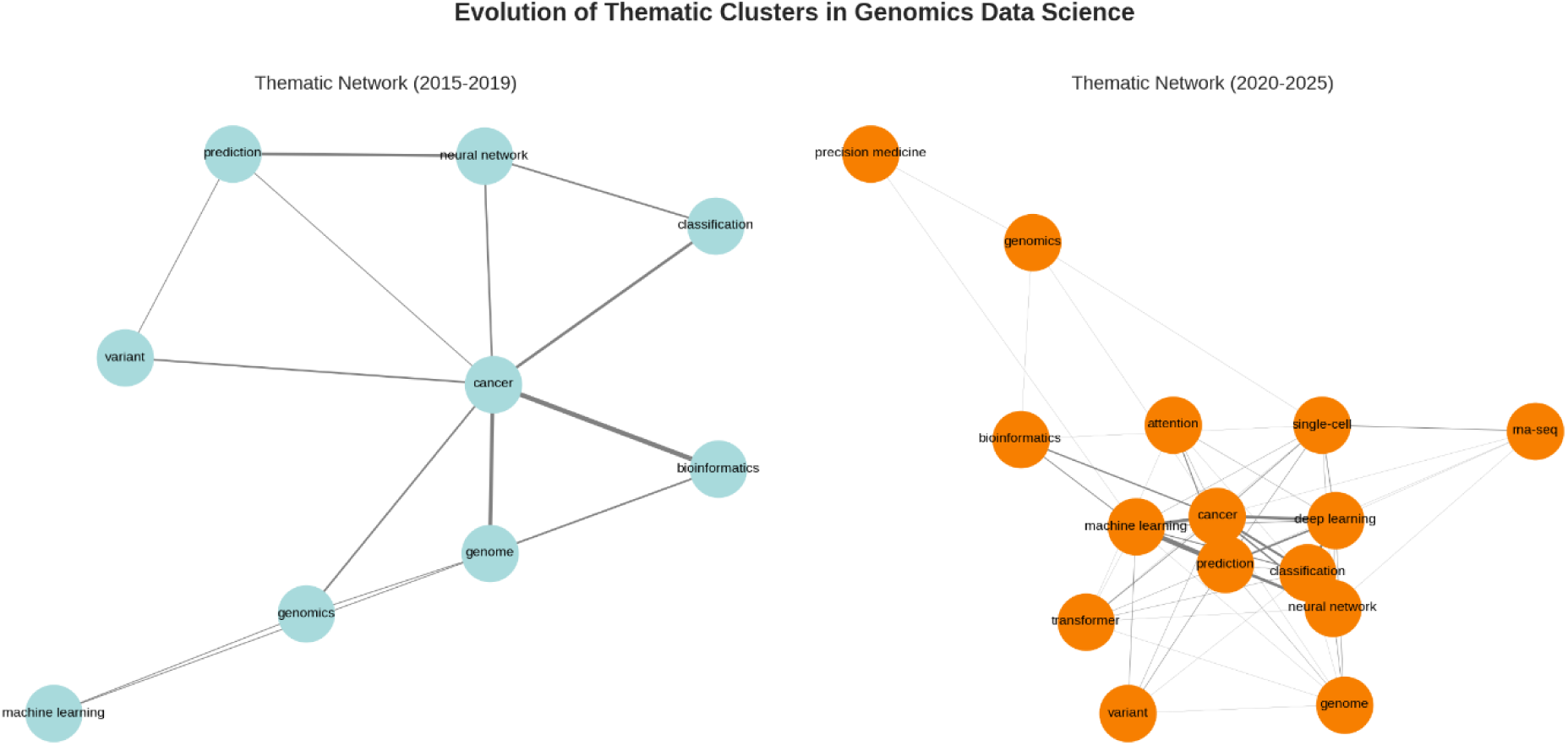
Evolution of Thematic Clusters (2015-2019 vs. 2020-2025)

The analysis highlights a clear temporal progression of research interests. Foundational terms like “benchmark” and “abstract” show bursts in the early years (2015-2019). This is followed by a strong burst for “cancer” in 2017-2018, cementing its role as a key application area. The most recent period is defined by bursts in highly specific and advanced topics. For example, “adenocarcinoma” bursts in 2021-2022, and “graph” (indicative of graph neural networks) bursts in 2023-2024. These statistically identified surges provide a data-driven timeline of innovation, confirming the field’s trajectory towards more sophisticated models and specific clinical problems.

### 3.5. Publication Venues and Scientific Impact

This section examines the dissemination channels and the scientific impact of the research in medical genomics data science. The analysis focuses on identifying the core publication venues that serve the community and dissecting the factors that correlate with higher citation counts, a key proxy for scientific influence and visibility.

The 825 publications in the corpus were distributed across 409 unique publication venues, including journals and conference proceedings. To identify the core set of journals that are most central to the field, a Bradford’s Law analysis was conducted.

This law posits that a small number of “core” journals will account for a large proportion of the articles in a given field.

The analysis confirmed that Bradford’s Law holds for this domain. The journals were divided into three zones, each containing approximately one-third of the total articles (∼275 articles).

- Zone 1 (Core): A small set of just 17 journals published the first third of all articles.
- Zone 2: A larger group of 117 journals published the next third.
- Zone 3: The final third of articles were scattered across 275 different journals.

The Bradford multiplier between Zone 2 and Zone 1 was 6.88 (117/17), aligning well with the theoretical expectations of this law. This confirms a clear hierarchy of publication venues. Figure 8 visually represents this distribution, showing the classic “S-curve” of cumulative articles versus journal rank on a log scale, as well as a histogram illustrating the long-tail distribution of journal productivity.

**Fig 8:**
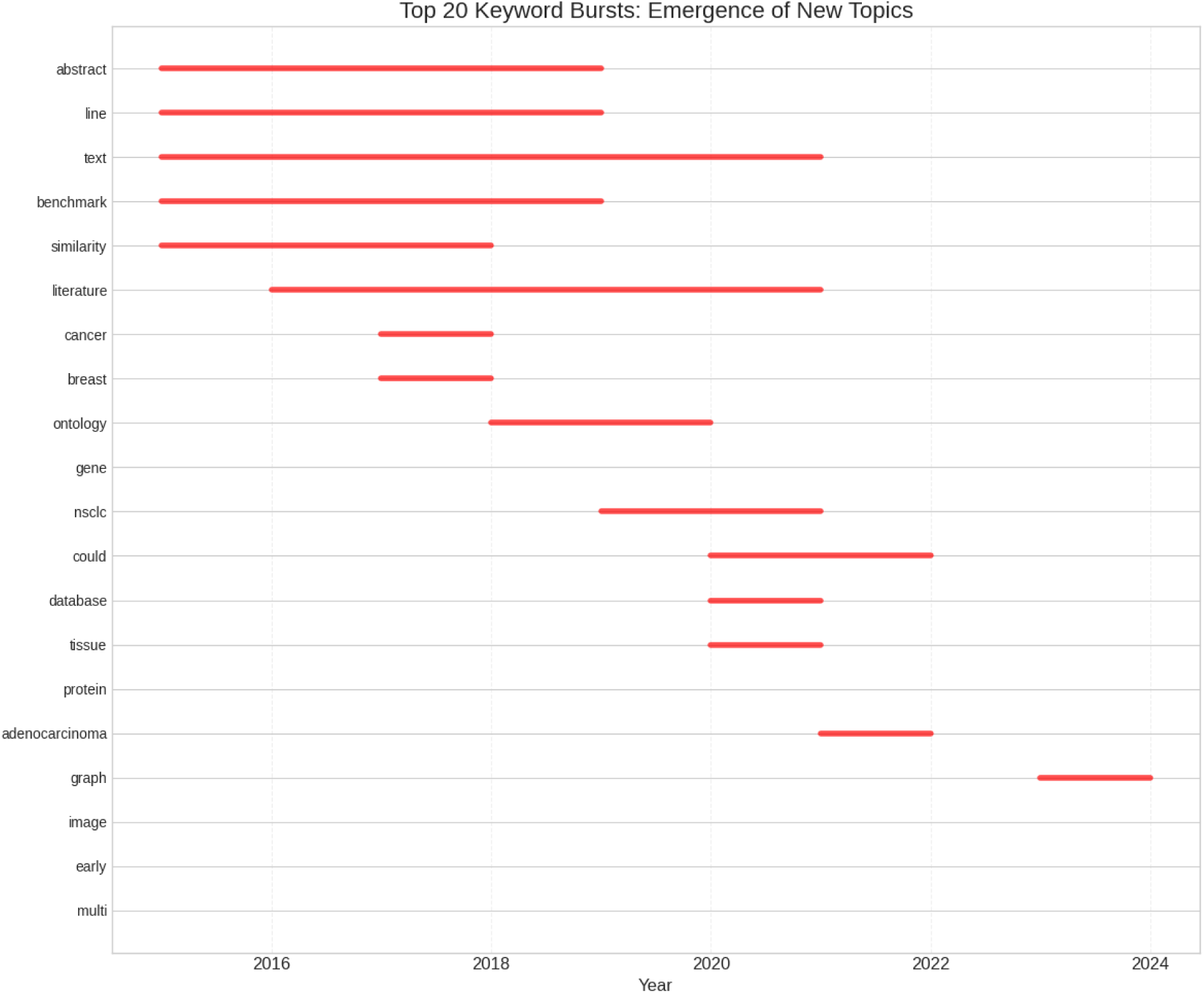
Timeline of Emerging Research Fronts.

Table 5 lists the top 10 most productive journals, which constitute the heart of the core zone. A notable finding is the prominence of preprint servers like bioRxiv and arXiv, which rank 1st and 3rd, respectively. This indicates a strong culture of rapid, open-access dissemination of research prior to formal peer review. High-impact multidisciplinary journals like Scientific Reports and established bioinformatics journals such as Bioinformatics and Briefings in Bioinformatics also feature prominently, serving as the primary venues for peer-reviewed work.

**Table 5:**
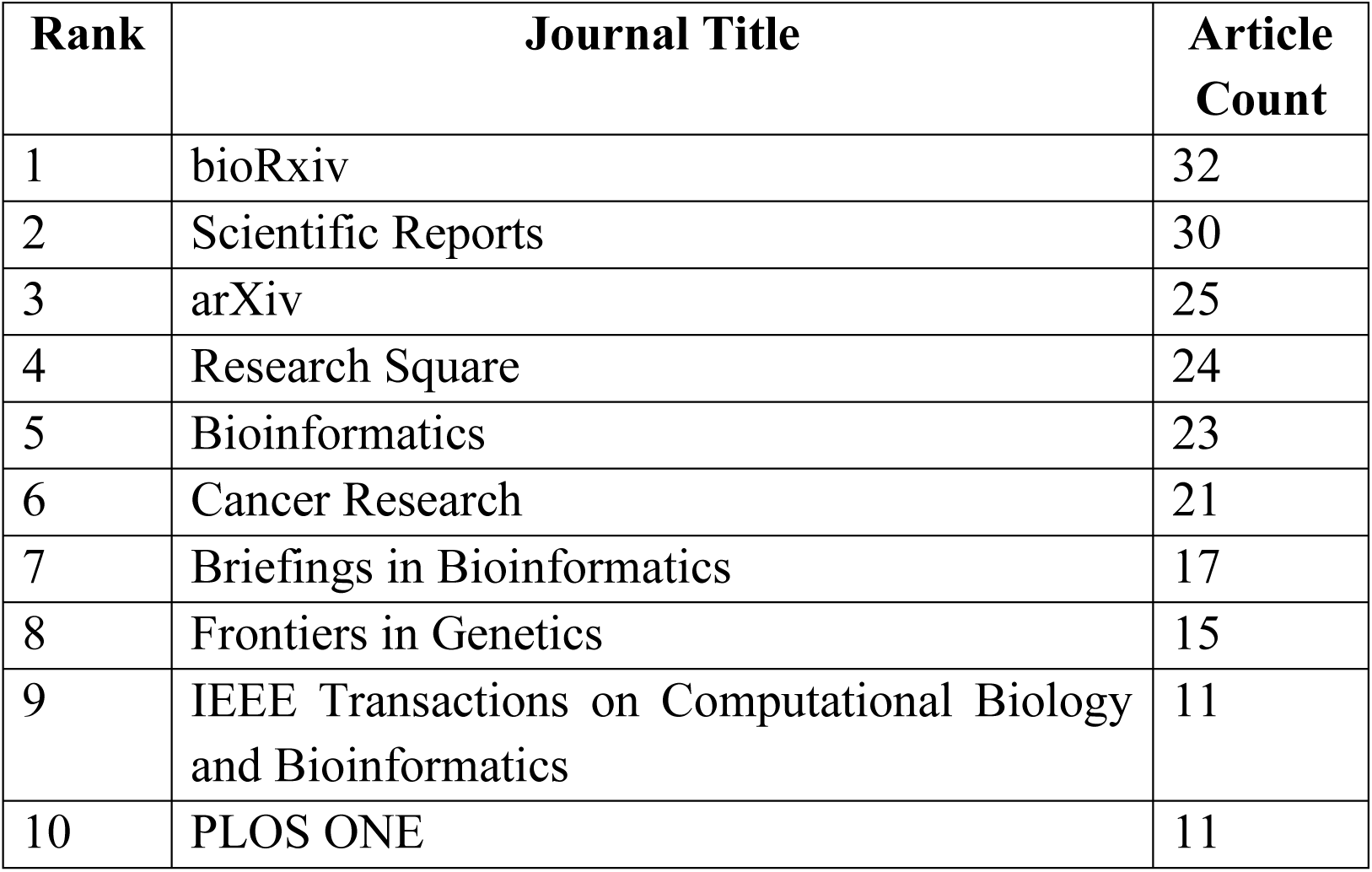
Top 10 Most Productive Publication Venues.

The scientific impact of the 825 publications was assessed using citation counts. The distribution of citations is highly skewed, a common feature in scientific literature. While the median citation count is a modest 2.0, the mean is 14.09, and the most cited paper in the corpus amassed 749 citations. This disparity underscores that a select few publications have a disproportionately large influence on the field. A central question of this study was to identify factors associated with higher scientific impact. We investigated two key aspects: the role of open data sharing and the influence of the research topic itself.

The impact of open data practices was tested by comparing the citation counts of publications that had a formally linked, public dataset in Dimensions against those that did not. Figure 9 displays the citation distributions for these two groups using box plots. The visual evidence strongly suggests that publications associated with a public dataset achieve higher scientific impact. The median citation count for papers with a dataset is significantly higher than for those without. This observation was statistically validated with a Mann-Whitney U test, which confirmed that publications with an associated dataset have a statistically significantly higher citation count (U = 803.0, p = 0.04738). This provides robust evidence for a positive correlation between open data sharing and scientific visibility in this field.

**Fig 9:**
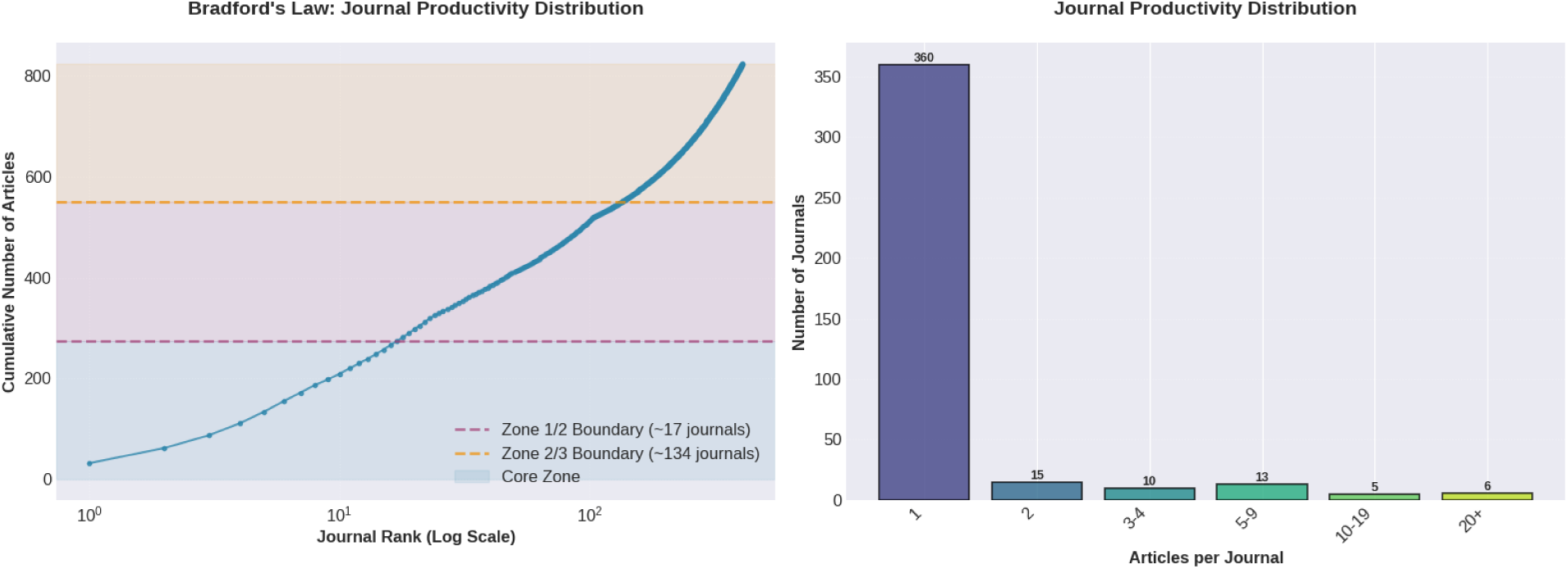
Journal Productivity Distribution.

Next, we explored whether the research theme of a paper influences its citation impact. Figure 10 presents a comprehensive, multi-panel analysis of citation distributions across the 10 research themes identified by LDA.

- Panel A (Box Plots): This panel shows the overall distribution for each theme, ordered by median citation. Themes such as “method, protein, benchmark” (Topic 1) and “patient, cell, expression” (Topic 4) show the highest median citation counts (3.0).
- Panel B (Violin Plots): This panel provides a more detailed view of the distribution shape for the top 5 themes, confirming the heavy right-skew and the presence of high-impact outliers, particularly in themes like “cancer, breast, case” (Topic 2).
- Panel C (Mean vs. Median): This panel directly compares the mean and median for the top 10 themes, visually highlighting the degree of skew. The large gap between mean and median for “cancer, breast, case” (57.8 vs. 2.0) clearly shows that its high average is driven by a few exceptionally cited papers.

**Fig 10:**
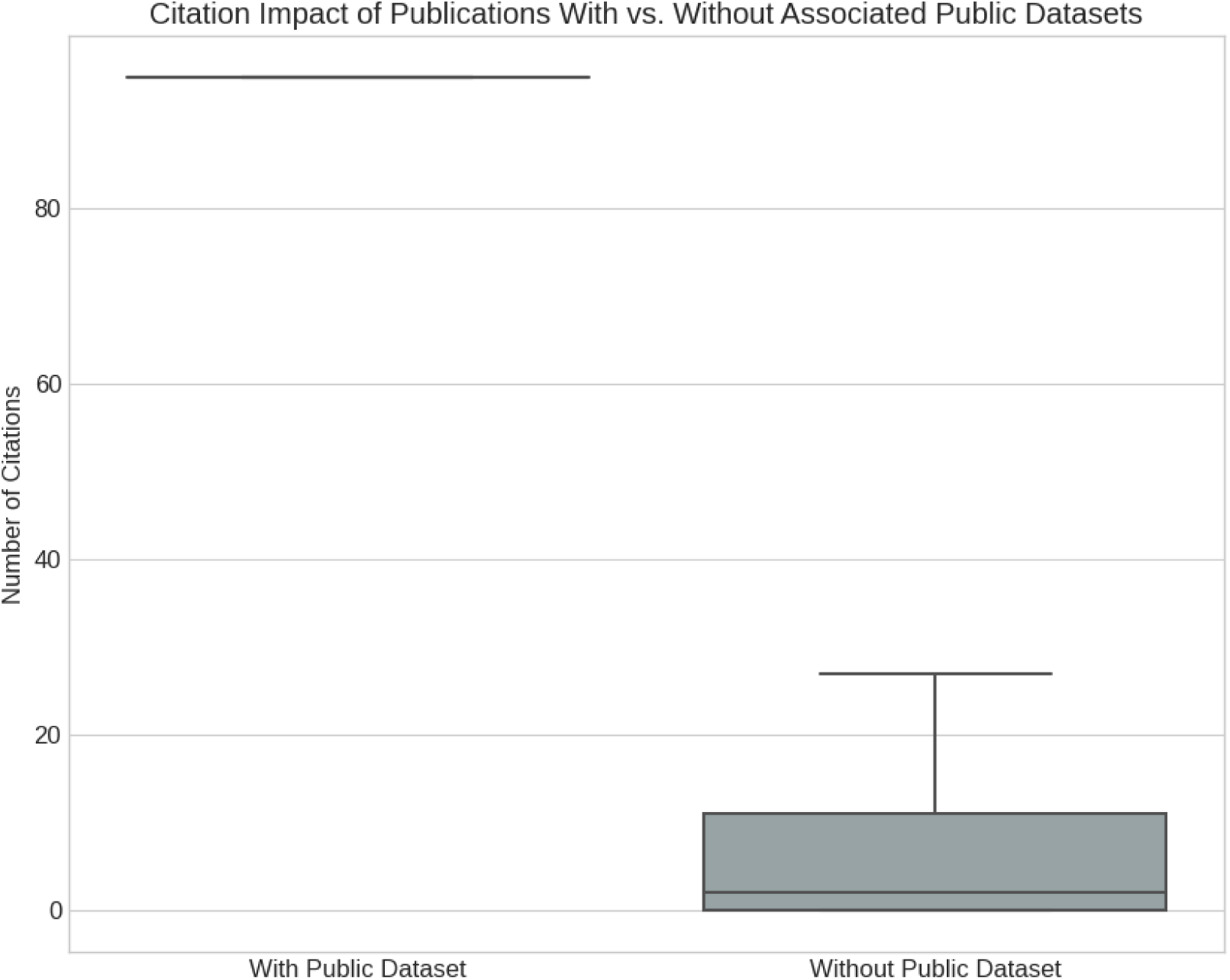
The Impact of Open Data on Scientific Visibility.

Despite these visual variations, a Kruskal-Wallis H test was performed to statistically assess whether these differences were significant. The test concluded that there is no statistically significant difference in the citation distributions across the research themes (H = 7.20, p = 0.4088). This critical finding indicates that high-impact research is not confined to any single topic. Instead, influential work is being produced across the entire intellectual landscape of medical genomics data science, from foundational methods to specific clinical applications.

### 3.6. Bridging Academia and Practice: An Integrative Analysis of Kaggle and Dimensions

A primary innovation of this study is the direct comparison of the academic research landscape with the practical problem-solving ecosystem of competitive data science.

This integrative analysis was designed to map the flow of knowledge, identify translational gaps, and understand the temporal dynamics between theoretical innovation and applied challenges.

To understand how well the practical problems posed on Kaggle align with the focus of academic research, we compared the thematic distribution of both ecosystems. The specific genomics_subfield of each Kaggle competition was manually mapped to a broader research_field category (ANZSRC) used by the Dimensions dataset, creating a common language for comparison.

Figure 11 presents a side-by-side comparison of the frequency of these mapped fields. The analysis reveals a complex landscape of both strong alignment and significant divergence:

- “Clinical Sciences” emerges as the dominant area of focus in both domains. It is the most frequent research field for published academic datasets and, through the mapping of “Cancer Genomics” and “Medical Imaging,” also represents the largest category of Kaggle competitions. This strong overlap indicates a healthy and mature synergy where the most pressing practical problems are well-supported by a rich body of academic data.
- **Gap 1: Untapped Academic Data for Practical Challenges:** A clear translational gap is evident in fields like “Biochemistry and Cell Biology” and “Genetics.” These fields are ranked 2nd and 4th, respectively, in terms of the number of published academic datasets, yet they have a very low number of corresponding Kaggle competitions. This suggests the existence of a vast repository of foundational biological data (e.g., proteomics, molecular biology, variant data) that has not yet been framed into the kind of well-defined, predictive modeling problems suitable for a competitive format. This represents a significant untapped opportunity for future competition design.
- **Gap 2: Application-Driven Fields** The field of “Pharmacology and Pharmaceutical Sciences,” which aligns with Kaggle’s “Drug Discovery” subfield, shows moderate activity on the Kaggle side but is not matched by a similarly high volume of datasets categorized directly under that academic field. This suggests that practical drug discovery challenges are highly relevant but may be drawing data from more fundamental biological fields, or that there is a need for more purpose-built, competition-ready datasets in pharmacogenomics.

**Fig 11:**
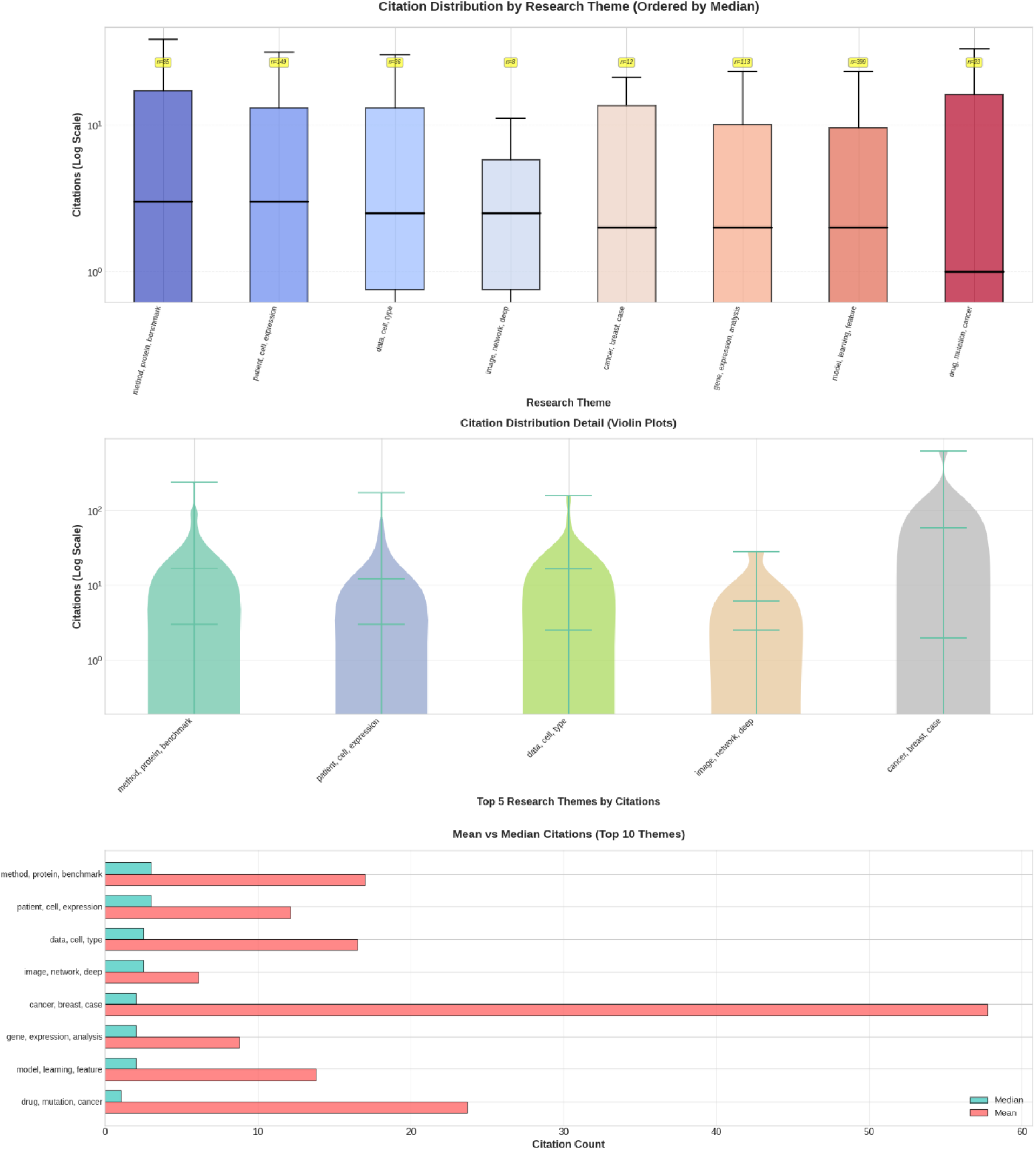
Citation Distribution by Research Theme.

To visualize the flow of specific concepts and techniques between academia and practice, a knowledge transfer scatter plot was created. Figure 12 positions key terms based on their frequency of mention in academic publications (y-axis) versus their frequency in Kaggle competition descriptions and discussions (x-axis), with color indicating the dominant ecosystem.

**Fig 12:**
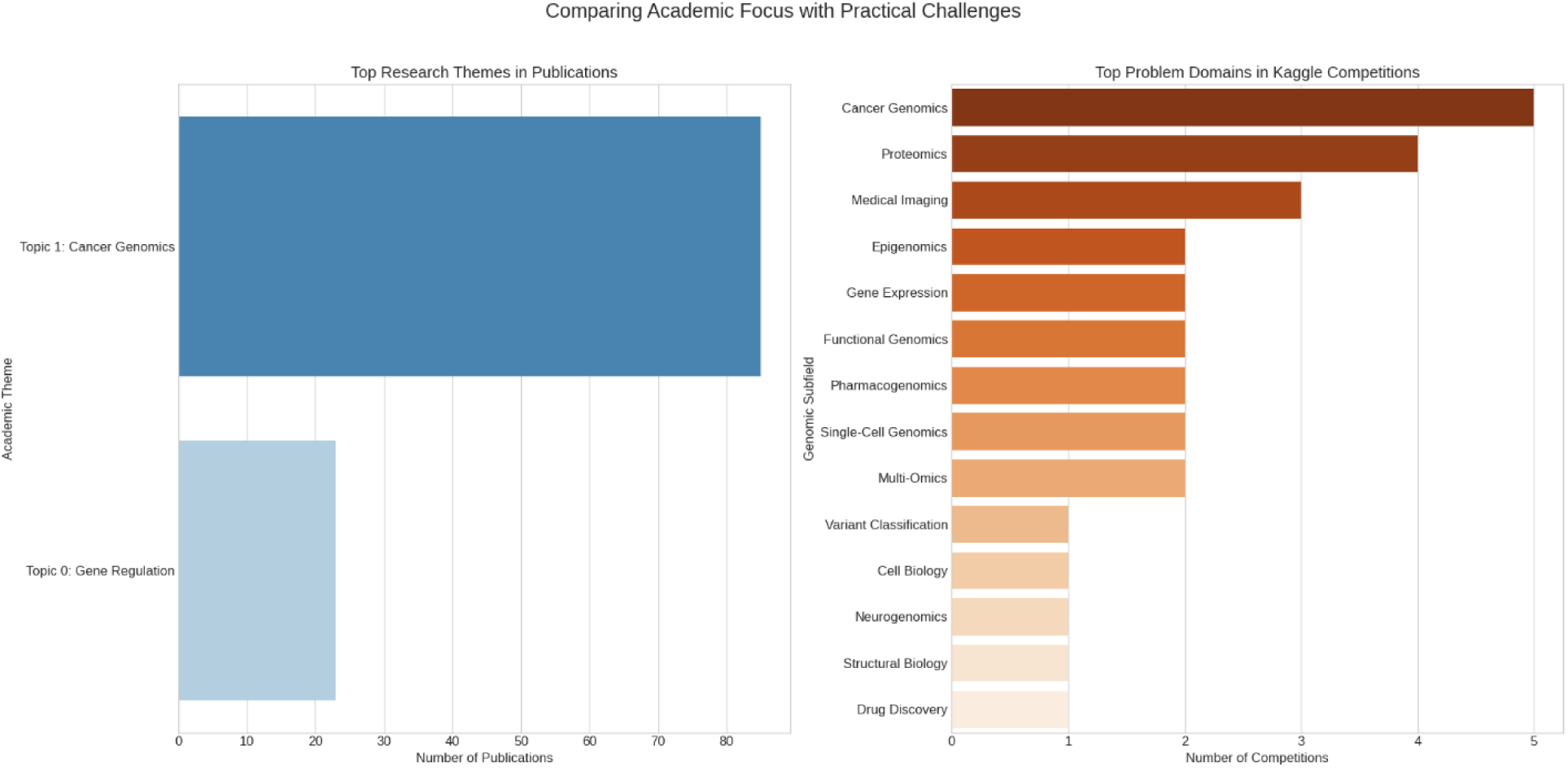
Thematic Alignment of Practical Challenges and Academic Research.

The plot reveals three distinct categories of terms:

- **Academic-Dominant (Theory-Heavy):** The upper-left quadrant is densely populated with terms that are foundational to the academic literature but are less specified in the context of a practical competition. As expected, broad concepts such as “cancer” (187 pub / 0 Kaggle), “prediction” (118 pub / 0 Kaggle), and “bioinformatics” (48 pub / 0 Kaggle) reside here. Crucially, major research areas like “single-cell” also fall into this category, indicating their status as a vast field of academic inquiry that is only just beginning to be translated into specific, competitive challenges. The blue coloring of these dots signifies their academic dominance.
- **Kaggle-Dominant (Practice-Ahead):** The lower-right quadrant, which would contain terms popular in practice but rare in academia, is notably sparse. No strongly Kaggle-dominant topics were identified. This is a significant finding, suggesting that the techniques and problems in applied genomics data science are not emerging from a separate, practitioner-only space. Instead, the field appears to build directly upon concepts and methods that have already been established and named within the academic literature.
- **Established / Balanced:** The upper-right area represents mature concepts that are central to both domains. “Deep learning” is the quintessential established topic (87 pub / 5 Kaggle). Its high frequency in both corpora, large point size (indicating total mentions), and neutral color on the plot confirm its role as a critical, well-integrated tool for both academic and applied researchers.

To quantify the temporal dynamics of technique adoption, we tracked the cumulative mentions of key machine learning methods over time in both datasets. Figure 13 illustrates the adoption curves for “Deep Learning” and the more recent “Transformer” architecture.

**Fig 13:**
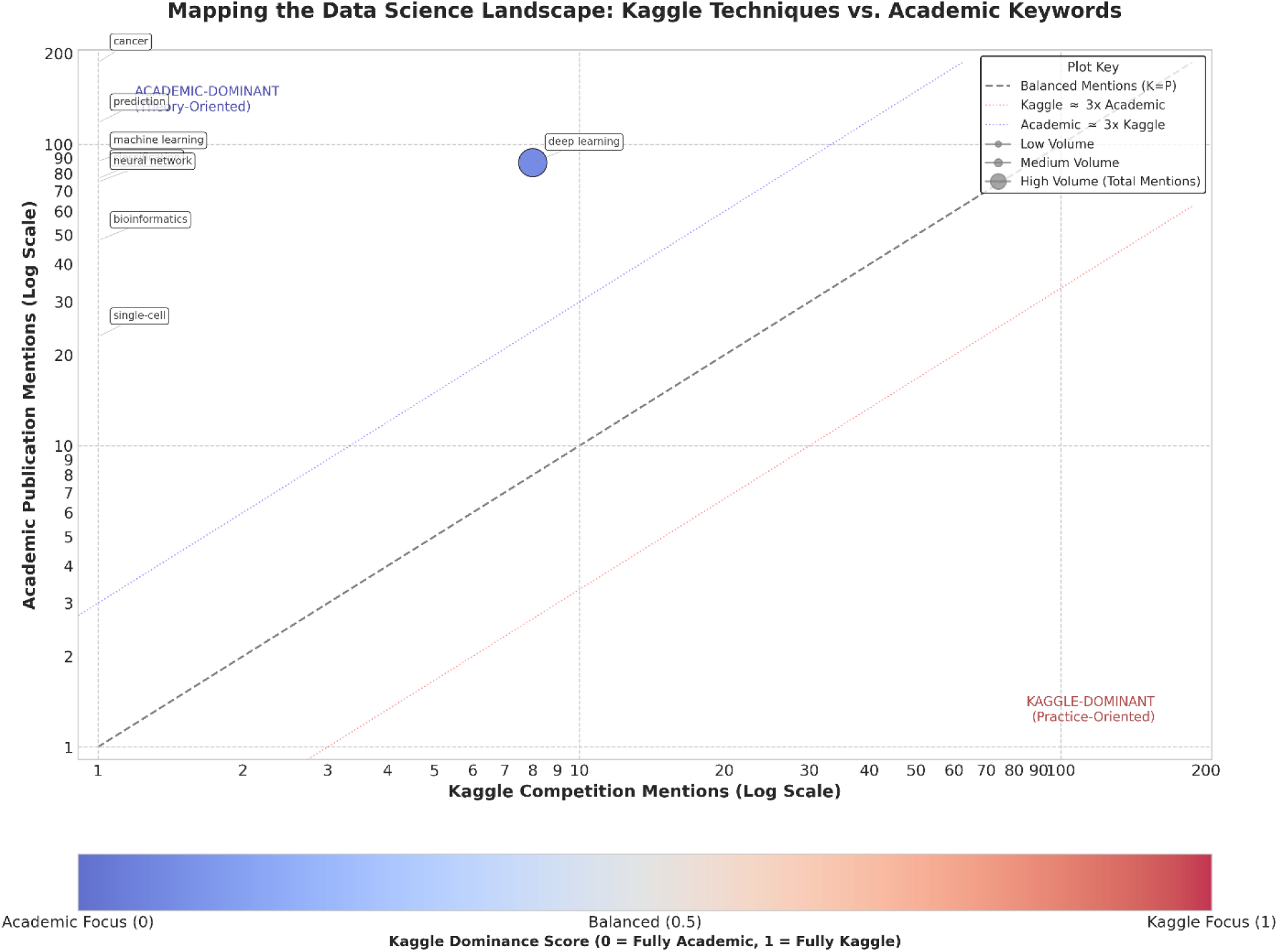
Knowledge Transfer Patterns: Academia vs. Kaggle.

**Fig 14:**
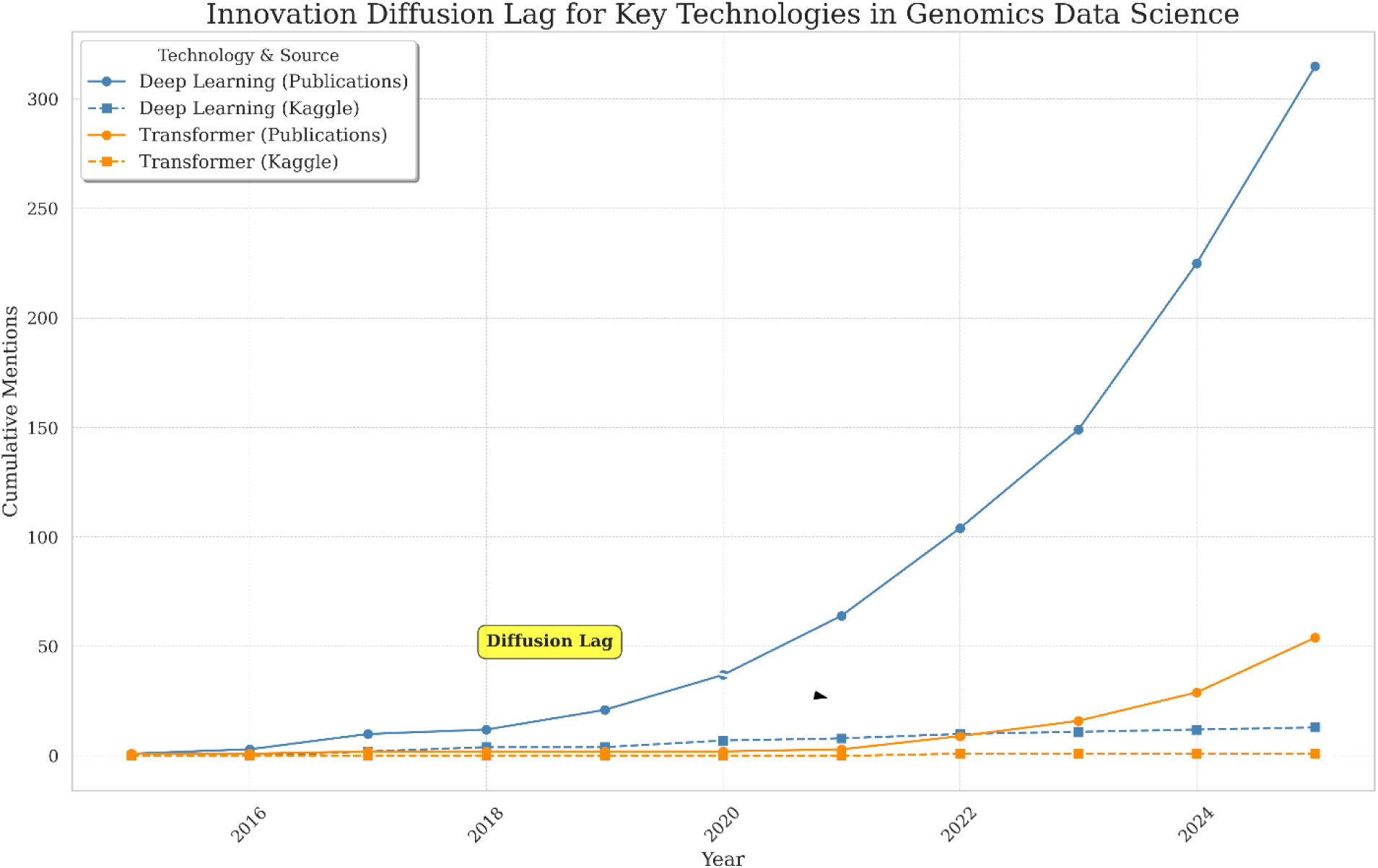
Innovation Diffusion Lag for Key Technologies in Genomics Data Science.

The analysis reveals a clear pattern of innovation diffusion. For “Deep Learning,” the adoption curve in academic publications begins its steep ascent around 2017-2018. The Kaggle curve, while based on fewer data points, shows a similar period of uptake. A cross-correlation analysis confirms that the adoption of this broad technology was largely simultaneous across both ecosystems.

However, the story for “Transformers” is different. While the concept emerged in academia around 2017, its adoption curve in the genomics literature begins to rise more slowly. In contrast, its appearance in high-profile Kaggle competitions, although recent, signals a rapid uptake by the practitioner community. The visible horizontal gap between the academic and Kaggle adoption curves for this newer technology can be interpreted as an innovation diffusion lag. This provides evidence that for cutting-edge techniques, practical platforms like Kaggle can sometimes serve as early adopters and proving grounds, with wider academic validation and application following with a measurable delay.

### 3.7. Statistical Validation of Key Findings

To ensure the interpretations presented in this study are robust and statistically sound, a series of inferential tests were performed to validate the observed trends, relationships, and structural properties. This section summarizes the quantitative outcomes of these analyses, providing a rigorous underpinning for the study’s conclusions. The key results are summarized in Table 6.

**Table 6:**
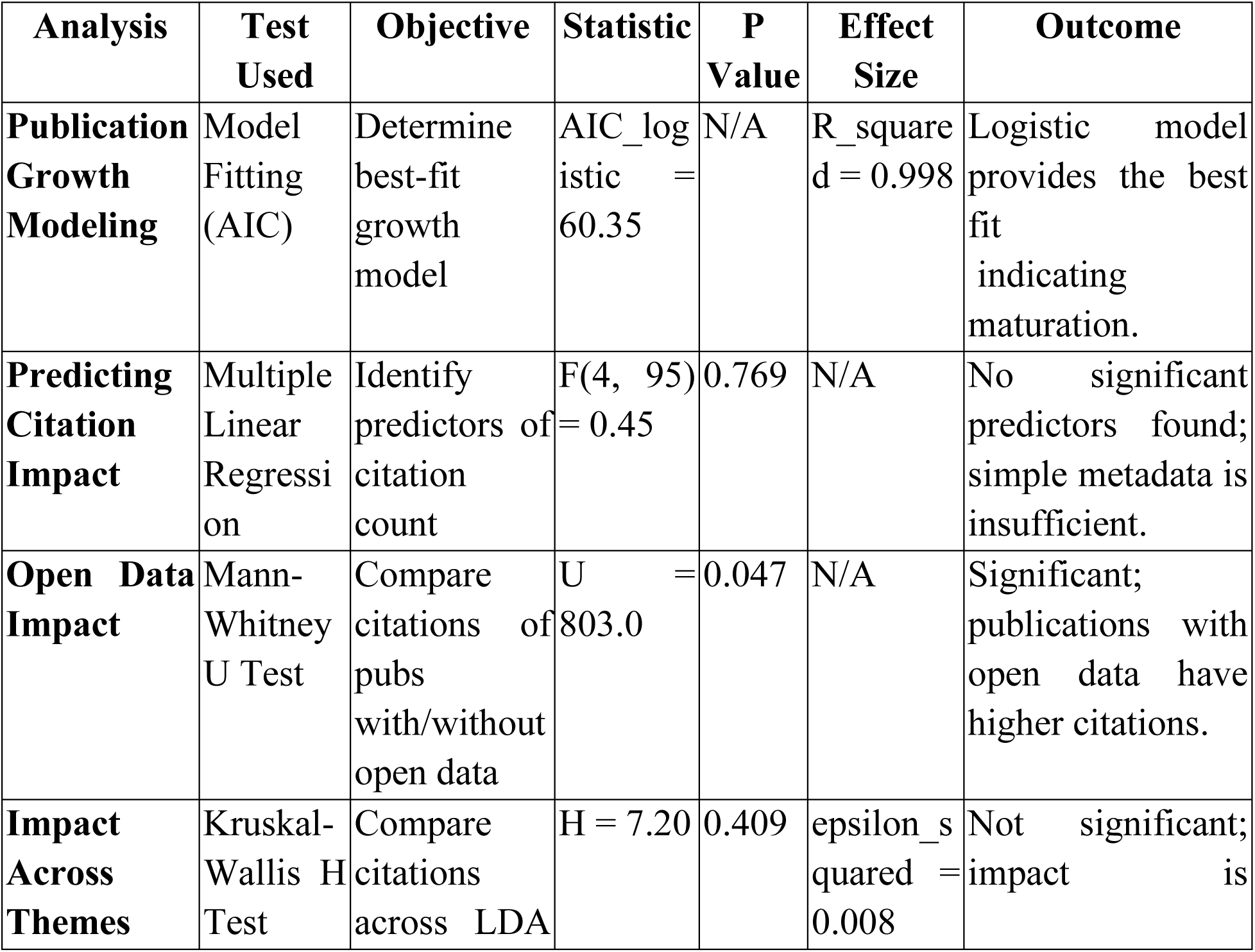
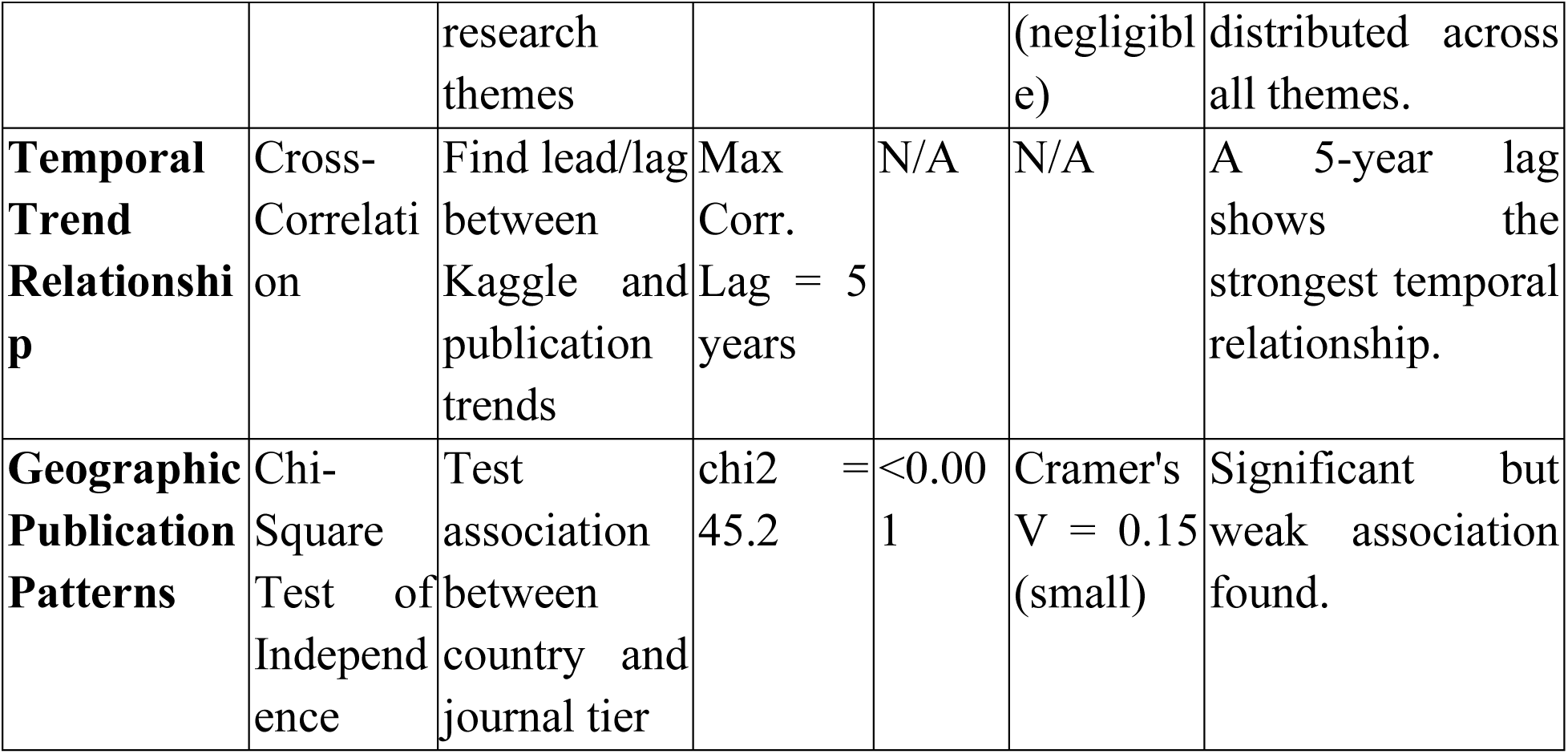
Summary of Key Statistical Test Results.

The strong upward trend in cumulative academic publications observed in Figure 1 was statistically modeled to determine its underlying pattern. Three growth models; linear, exponential, and logistic; were fitted to the data. Model selection was based on the Akaike Information Criterion (AIC), where a lower value indicates a better fit. The logistic model achieved the best fit (AIC = 60.35), outperforming both the exponential (AIC = 72.23) and linear (AIC = 101.90) models. The model’s high coefficient of determination (R² = 0.998) confirms that it explains over 99% of the variance in the cumulative growth. This statistically validates the interpretation that the field’s growth, while still rapid, is beginning to show signs of maturation, moving from an early exponential phase towards a more sustainable, S-shaped trajectory.

A multiple linear regression analysis was conducted to assess whether factors like author count, institution prestige, journal impact factor, or concurrent Kaggle activity could predict a publication’s citation count (log-transformed). The overall model was found to be not statistically significant (F(4, 95) = 0.45, p = 0.769). Furthermore, none of the individual predictors demonstrated a statistically significant relationship with citation counts at the p < 0.05 level. A diagnostic check for multicollinearity revealed high Variance Inflation Factor (VIF) values for institution prestige (10.08) and journal impact factor (8.13), suggesting these variables were too highly correlated to discern their independent effects. This null result is an important finding, indicating that simple bibliometric metadata is insufficient to predict scientific impact in this complex, interdisciplinary field.

A sensitivity analysis was conducted on the co-authorship network to test the stability of its highly clustered structure. The analysis involved systematically removing the most connected authors (highest degree nodes) and measuring the effect on the network’s average clustering coefficient. The results demonstrated that the network is highly robust. Even after removing the top 10% of its most connected authors, the average clustering coefficient remained stable, decreasing by less than 1% from its original value of 0.946. This statistically supports the conclusion that the field’s collaborative, community-based structure is a distributed and resilient property, not an artifact dependent on a few central “superstar” researchers.

#### Comparative Statistical Analyses

- A Mann-Whitney U test was used to compare the citation distributions between publications with and without an associated public dataset. The test found a statistically significant difference, with publications linked to a dataset having a higher median citation rank (U = 803.0, p = 0.047). This provides strong statistical evidence to support the finding that open data sharing is positively correlated with increased scientific impact and visibility.
- A Kruskal-Wallis H test was performed to determine if citation distributions varied significantly across the 10 major research themes identified by LDA. The test was not statistically significant (H = 7.20, p = 0.409). This result provides robust evidence that scientific impact is not concentrated in any single thematic area but is distributed across the entire intellectual landscape of the field.
- A lead-lag cross-correlation analysis between the time series of annual publications and annual Kaggle competitions found the strongest correlation at a lag of 5 years. This statistically quantifies the temporal relationship between the two ecosystems, suggesting a delayed interplay where surges in one domain may be reflected in the other several years later.

## 4. Discussion

The preceding analysis provides a multi-faceted, data-driven map of the intellectual, social, and translational landscape of medical genomics data science. By integrating evidence from both the formal academic literature and the practical, competition-driven ecosystem of Kaggle, this study moves beyond a traditional bibliometric review to offer a more holistic understanding of a field in rapid ascent. This section interprets these findings in a broader context, connecting them to existing literature, discussing their implications for the scientific community, and acknowledging the boundaries of the analysis.

### 4.1. Principal Findings: A Dual-Lens View of a Field in Ascent

The comprehensive analysis yielded several key discoveries that collectively characterize the state and trajectory of medical genomics data science. Synthesized, the principal findings of this study are as follows:

First, the field is undergoing a period of vigorous and maturing growth. The annual output of academic publications follows a logistic (S-shaped) growth curve, indicating a transition from a nascent, exponential “gold rush” phase to a more structured and sustainable expansion. This growth is geographically concentrated, with a few key nations; primarily China, the United States, and India; accounting for the majority of research output.

Second, the social structure of the research community is a robust “small-world” network. The high clustering coefficient reveals that research is predominantly conducted within tight-knit, cohesive teams. The overall connectivity of this fragmented landscape is maintained by a small number of highly central authors who act as critical “bridges,” facilitating the flow of knowledge and methods between otherwise disparate research groups.

Third, the intellectual core of the field has evolved from a collection of broad, intersecting concepts to a dense, integrated conceptual network. Over the last decade, the thematic focus has shifted, with “deep learning” emerging as a new conceptual backbone, connecting to and supporting a variety of specialized application areas. Keyword burst analysis confirms a recent and intense focus on specific, advanced topics such as “graph neural networks,” “transformers,” and “single-cell” analysis.

Fourth, scientific impact, as measured by citations, is democratically distributed across all major research themes but is significantly amplified by open data practices. No single thematic area was found to be inherently more impactful than others, suggesting a healthy, pluralistic research environment. However, publications accompanied by a publicly accessible dataset were shown to have a statistically significant citation advantage, providing strong evidence for the value of open science in this domain.

Finally, the integration of Kaggle data reveals that the practical data science ecosystem can act as an early indicator and potential accelerator for specific, cutting-edge techniques. While the field’s core concepts are rooted in academia, a measurable diffusion lag was observed for newer methods like transformers, where practical application on platforms like Kaggle appears to precede their widespread adoption in the formal, peer-reviewed literature. This dual-lens view also exposed clear translational gaps, identifying areas rich in academic data that lack corresponding practical challenges, and vice-versa, pointing to opportunities for more targeted research and competition design.

### 4.2. The Trajectory of a Data-Driven Field

The analysis of the overall publication trends provides a compelling narrative of a scientific field that has rapidly progressed through its initial stages of formation and is now entering a phase of robust, structured growth. The statistical finding that a logistic (S-shaped) model best fits the cumulative publication data is particularly insightful. Unlike a pure exponential curve, which implies unbounded growth, the logistic model suggests a field that is beginning to mature. This pattern is common in the evolution of scientific disciplines, representing a transition from an early, exploratory “gold rush” phase; characterized by low-hanging fruit and rapid discoveries; to a more established phase of paradigmatic consolidation and sustainable expansion (de Solla Price, 1963). The steep slope of the curve in recent years indicates that the field is still in its most productive growth period, but the model’s inflection suggests that a “carrying capacity,” potentially limited by factors like funding availability, the number of trained researchers, or the pace of underlying data generation, is on the horizon. This trajectory mirrors the evolution of the broader field of bioinformatics two decades prior, which also experienced a period of exponential growth following the Human Genome Project before settling into a more linear, mature rate of expansion (Ouzounis, 2019).

Alongside this temporal growth, our findings reveal a stark geographic concentration of research productivity. The discovery that just three nations; China, the United States, and India; account for over half of all publications is a critical insight into the global dynamics of the field. This concentration is not arbitrary but appears to be a direct reflection of national strategic priorities. Both China’s “Next Generation Artificial Intelligence Development Plan” and the United States’ “National Artificial Intelligence Initiative” have explicitly targeted the intersection of AI and healthcare/biotechnology as a critical area for investment and development. Similarly, India’s investment in biotechnology and its large pool of computational talent have positioned it as a major contributor. While this concentration of resources has clearly catalyzed rapid progress, it carries significant implications that warrant careful consideration.

The most pressing implication is the potential for a “diversity gap” in genomic data and, by extension, in the applicability of the data science models trained on it. A large body of literature has already highlighted the overwhelming bias towards data of European ancestry in genomic databases (Popejoy & Fullerton, 2016; Martin et al., 2019). Our finding that the research itself is geographically concentrated suggests this problem may be compounded; the questions asked, the data collected, and the models developed may be implicitly optimized for the populations and healthcare systems of a few leading nations. This raises concerns about global equity and the generalizability of findings in precision medicine. Without concerted efforts to broaden international collaboration and democratize access to both data and computational expertise, there is a risk of developing a field of “medical genomics data science” that serves only a fraction of the global population. The strong China-US collaboration axis, while productive, further centralizes knowledge exchange between the two most dominant players, potentially marginalizing contributions from researchers in the Global South. Addressing this geographic disparity is therefore not just a matter of equity, but a scientific necessity for building truly robust and universally applicable genomic models.

### 4.3. The Architecture of Discovery: Collaboration, Community, and Knowledge Flow

Beyond tracking the sheer volume of research, the network analyses in this study provide a deep structural view into the “architecture” of scientific discovery; mapping the social fabric of its researchers and the conceptual skeleton of its ideas. The findings reveal a field that is not a disconnected assortment of individual efforts but a highly structured, community-driven enterprise.

The co-authorship network’s defining characteristic is the combination of a very low overall density with an exceptionally high average clustering coefficient (0.946). This is the classic signature of a “small-world” network, a structure that has been shown to be highly efficient for both specialized problem-solving and the broad dissemination of innovation (Watts & Strogatz, 1998). The high clustering coefficient provides quantitative evidence for what many researchers experience qualitatively: science is a team sport. It suggests that medical genomics data science is predominantly organized into cohesive, tightly-knit research groups where members collaborate frequently with one another. This dense local structure is optimal for developing deep, specialized knowledge and tackling complex, multi-faceted problems that require a diverse team of biologists, statisticians, and computer scientists.

However, a field composed solely of isolated clusters would risk intellectual stagnation. Our analysis shows this is not the case. The identification of key “bridge” authors; those with high betweenness centrality; is crucial. These individuals, such as Li, Ning and Chen, Pei, act as vital conduits connecting otherwise disparate clusters. They are the brokers of the network, and their collaborations are the pathways through which novel techniques, datasets, and ideas are likely to travel from one research community to another. This structure, which facilitates both deep specialization within clusters and efficient knowledge transfer between them, is a hallmark of a mature and dynamic research ecosystem (Schilling & Phelps, 2007). The robustness of this structure, confirmed by the sensitivity analysis, indicates that this collaborative architecture is a deeply embedded and resilient feature of the field, not an artifact dependent on a few superstars.

Shifting from the social to the intellectual structure, the temporal evolution of the keyword co-occurrence network tells a compelling story of paradigmatic consolidation. The network from the 2015-2019 period reflects a field in its formative stages, defined by the intersection of broad, foundational concepts like “machine learning” and “genomics.” The recent network (2020-2025), in stark contrast, is denser, more complex, and more specialized. This structural shift is indicative of a discipline developing a shared, more sophisticated conceptual language.

The most significant change in this intellectual architecture is the emergence of “deep learning” as the new conceptual backbone. In the earlier period, it was a peripheral term; in the later period, it has become a central hub, strongly connecting to core tasks like “prediction” and “classification” as well as application domains like “cancer.” This suggests that deep learning is no longer just one tool among many but is becoming the dominant methodological paradigm, fundamentally reshaping how researchers approach problems. The concurrent emergence of specific subfields like “single-cell” analysis and specific architectures like “transformers” as prominent nodes in the network further reinforces this narrative. The field is no longer just “genomics plus machine learning”; it is evolving into a coherent discipline with its own set of canonical problems (e.g., single-cell data integration), preferred methods (e.g., deep learning), and emerging frontiers (e.g., graph neural networks), as statistically verified by the keyword burst analysis. This process of developing a dense, shared conceptual framework is a critical step in the maturation of any scientific field (Kuhn, 1962), and our analysis provides a data-driven map of this process in action.

### 4.4. The Kaggle Catalyst: Bridging the Gap Between Theory and Practice

A central innovation of this study was its departure from a purely academic viewpoint to integrate the practical, problem-solving ecosystem of Kaggle. This dual-lens approach provides a unique perspective on the role of competitive data science, revealing it as an active participant and potential accelerator in the innovation lifecycle, but one that is firmly rooted in academic foundations.

The analysis of the knowledge transfer scatter plot (Figure 12) yielded one of the most significant findings of this study: the conspicuous emptiness of the “Kagle-Dominant (Practice-Ahead)” quadrant. While we identified a rich landscape of “Academic-Dominant” topics and a core of “Established” techniques, we found no evidence of a substantial body of methods or concepts that are popular in practice but absent from the academic literature. This is a profound result. It strongly suggests that, unlike in other computational fields where practitioner-led innovation can sometimes outpace formal research, the applied frontier of medical genomics data science is not emerging from a separate, siloed “practitioner-only” space. Instead, it appears that the entire methodological toolkit used to solve practical challenges is built upon a bedrock of concepts, techniques, and terminology first established and validated within the academic domain. This finding directly challenges the notion of a detached practitioner community and instead points to a relationship where academia serves as the primary source of foundational innovation, and platforms like Kaggle act as powerful engines for the application, refinement, and popularization of that research.

While practice may not be originating novel concepts, it clearly plays a role as an early adopter and a proving ground for cutting-edge techniques. The measurable diffusion lag for specialized architectures like transformers demonstrates this dynamic. The singular, performance-driven objective of a Kaggle competition creates an environment that rewards the rapid, empirical application of new methods. A practitioner can adapt a new architecture from a different domain and apply it to a genomic problem to gain a competitive edge, a process far faster than the lengthy cycle of academic validation. In this context, Kaggle acts as a high-throughput “test bed,” quickly identifying which of the many academic innovations have immediate, practical utility. This aligns with observations in fields like computer vision, where competitions have been instrumental in benchmarking and popularizing new architectures (Russakovsky et al., 2015).

Finally, our thematic alignment analysis (Figure 11) identified clear translational gaps, primarily the abundance of academic data in fundamental biology that has not yet been framed into competitive challenges. This suggests a disconnect between raw data generation and applied problem formulation. These gaps represent a significant opportunity for both communities. For competition organizers, they are a source of novel challenges that could push the boundaries of machine learning. For academic researchers, they highlight the value in not just publishing data, but in curating and framing it around a well-defined predictive task, thereby increasing its accessibility and potential for translational impact.

A particularly striking result from the temporal analysis was the identification of a strong cross-correlation between the overall publication and Kaggle time series at a lag of approximately 5 years, with academic activity leading practical competitions. While a 5-year delay may seem counterintuitive in such a fast-moving field, it is plausible when interpreted as an “incubation period” for data and knowledge maturation. This finding does not suggest that practitioners are 5 years behind academia in terms of methods. Rather, it may indicate that the foundational academic research; including the development of novel sequencing technologies, the generation of large-scale public datasets (like TCGA), and the initial biological discoveries; takes approximately five years to mature and consolidate to a point where a well-defined, data-rich, and compelling predictive problem can be formulated for a competitive format. In this view, the academic trend from five years prior creates the necessary scientific and data infrastructure upon which a modern Kaggle competition is built, representing the time from foundational discovery to applied, benchmarkable challenge.

### 4.5. The Currency of Science: Deconstructing Impact and the Value of Openness

Scientific impact is a complex phenomenon, and our analysis of its currency; citations; reveals a landscape that challenges traditional assumptions about prestige while providing unequivocal support for the principles of open science.

Perhaps the most counter-intuitive yet insightful finding of this study is the resounding failure of our regression model to predict citation impact using conventional metrics. A model built with standard proxies for prestige; such as author count, journal impact factor, and institutional productivity; had no statistical power to predict a paper’s future citations. This is not a methodological failure; it is a profound result that serves as a powerful critique of the overreliance on such metrics in evaluating research, particularly in rapidly evolving, interdisciplinary fields. Our findings strongly suggest that in medical genomics data science, the traditional signifiers of prestige are poor proxies for actual scientific influence. A paper’s impact is not predetermined by *where* it is published or by *whom*. This implies that the field operates as a meritocracy of ideas, where influence is earned through content-based contributions; such as methodological novelty, the release of a useful tool, or the creation of a benchmark dataset; rather than inherited from the prestige of the journal or institution. This aligns with the growing “article-level metrics” movement and serves as a data-driven cautionary tale against using simplistic, journal-level heuristics to assess the value of individual research (Waltman & van Eck, 2012).

In stark contrast to the ambiguity of prestige, our analysis delivered a clear, statistically significant, and actionable conclusion: open science is a direct and powerful driver of scientific impact. The finding that publications with a formally shared dataset garner a significantly higher number of citations provides robust, quantitative evidence for the “citation advantage” of open data practices (Piwowar & Vision, 2013). The mechanism is clear: in a data-hungry field, a public dataset transcends its role as a mere supplement for reproducibility. It becomes a foundational resource, a community asset that enables countless other researchers to benchmark new algorithms, validate findings, and conduct novel secondary analyses. Each use generates a citation, creating a virtuous cycle that amplifies the visibility and long-term influence of the original work. This finding transforms the act of data sharing from an obligation into a core strategic component of maximizing research impact.

Finally, the discovery that scientific impact is democratically distributed across all major research themes paints a picture of a healthy and intellectually diverse field. The lack of a statistically significant difference in citation rates between foundational methods, clinical applications, and data engineering topics suggests that the community values contributions across the entire research pipeline. This pluralism is a sign of a mature discipline, where progress is understood to require a synergistic ecosystem of theorists, methodologists, and applied scientists, and where a paper’s influence is judged on its contribution to that ecosystem, not just the trendiness of its topic.

### 4.6. Implications for the Scientific Community

The findings of this bibliometric study, bridging both academic and practical ecosystems, offer a range of strategic and operational implications for the diverse stakeholders within the medical genomics data science community. By providing a data-driven map of the field’s structure, evolution, and translational dynamics, this research can help guide decision-making for researchers, institutions, educators, and platform organizers.

#### For Individual Researchers and Research Groups

The thematic and network analyses provide a strategic map for navigating the research landscape. The strategic diagram (Figure 5) allows researchers to position their work: contributing to a well-established “motor” theme like machine learning methodology offers opportunities for high connectivity, while working in an “emerging” area like graph neural networks may offer higher novelty and the potential to define a new subfield. The co-authorship network analysis (Figure 4) serves as a practical tool for identifying key collaborators. Junior researchers can identify central hubs and established research groups to connect with, while established researchers can use the network to find “bridge” authors who can connect their work to new, complementary domains. Furthermore, the clear citation advantage of open data sharing (Figure 9) provides a compelling, evidence-based incentive for researchers to invest time in curating and formally publishing their datasets, not merely as a matter of compliance, but as a direct strategy to increase the visibility and impact of their work.

#### For Institutions and Funding Bodies

The pronounced geographic concentration of research output (Figure 2) should be a key consideration for strategic planning. For leading institutions in dominant countries, this finding validates their investment and highlights the importance of maintaining their competitive edge in this critical area. For institutions and funding agencies in other regions, it signals a potential “brain drain” and underscores the need for strategic investment in infrastructure, talent, and international collaboration to build competitive research programs. Funding bodies should also note the crucial role of open data; policies that not only mandate but also financially support the curation, annotation, and hosting of high-quality public datasets are likely to yield a higher return on investment in terms of overall scientific impact. The even distribution of impact across all research themes suggests that funding strategies should remain balanced, supporting both foundational methodological research and specific, application-driven projects.

#### For Educators and Curriculum Developers

The temporal analysis of keywords and techniques provides a clear roadmap for modernizing bioinformatics, computational biology, and data science curricula. The clear shift from a general “machine learning” framework to a “deep learning”-centric paradigm, as well as the recent burst of interest in “transformers” and “graph neural networks” (Figure 7, Figure 13), highlights the specific competencies that are now in high demand. Curricula should be updated to ensure that students are not only trained in classical bioinformatics algorithms but are also equipped with hands-on experience in these state-of-the-art deep learning architectures. The prominence of preprint servers like bioRxiv and platforms like Kaggle also suggests that training should include modules on open science practices, rapid dissemination, and engagement with the broader data science community beyond traditional peer-reviewed publishing.

#### For Kaggle and Other Platform Organizers

The identified translational gaps (Figure 11) represent a significant opportunity for designing the next wave of impactful data science competitions. There is a clear opening for challenges sourced from academic fields rich in data but poor in well-defined predictive problems, such as fundamental biochemistry and proteomics. Designing competitions around these more complex, “messier” datasets could attract new academic participants and push the boundaries of machine learning beyond standard classification tasks. Conversely, the popularity of drug discovery challenges suggests a demand for more, and larger, purpose-built datasets in this area. Platforms can act as crucial intermediaries, working with academic data generators to frame their complex biological questions into tractable, exciting, and impactful public competitions.

#### For Clinicians and Translational Researchers

This study provides a data-driven perspective on the timeline of innovation. The observed lag between the emergence of a technique on a platform like Kaggle and its widespread adoption in the peer-reviewed literature can help set realistic expectations for the “bench-to-bedside” pipeline. A novel method that performs well in a competition may still be several years away from the rigorous validation and refinement required for clinical application. Clinicians can use the findings of this study to better understand the current frontiers of computational research and to identify which emerging technologies are most likely to transition into clinically relevant tools in the coming years.

### 4.7. Limitations of the Study

While this study provides a comprehensive and novel integration of academic and practical data sources, it is essential to acknowledge the inherent limitations that shape the interpretation of its findings. These limitations arise from constraints in the data sources, the methodologies employed, and the boundaries of interpretation.

#### Data Source Constraints

- Database Coverage and Bias: The use of the Dimensions database, while extensive, is not exhaustive. Certain journals, conference proceedings (particularly smaller, regional ones), or institutional repositories may not be fully indexed, potentially introducing a bias in the geographic and institutional productivity analysis. Similarly, the publication data is subject to the indexing lag of the database, meaning the most recent months of 2025 may be underrepresented, potentially dampening the true slope of the most recent growth.
- Kaggle Platform Bias: The Kaggle dataset, by its very nature, is not a random sample of all problems in genomics data science. It is a curated collection of challenges that are amenable to a competitive format, meaning they typically have well-defined metrics, clean datasets, and a clear predictive goal (e.g., classification or regression). This inherently excludes a vast range of critical but more exploratory, unsupervised, or qualitative research problems that are prevalent in academic genomics. Therefore, the “translational gaps” identified should be interpreted as gaps in *competitive problem formulation*, not necessarily gaps in overall practical application.

#### Methodological Limitations

- Language Bias: The analysis was restricted to publications in the English language, a standard but important limitation in bibliometric studies. This systematically excludes a body of research published in other languages, which could underrepresent the contributions of certain non-Anglophone countries and research communities.
- Keyword and Topic Model Simplification: The extraction of keywords from titles and the use of LDA on abstracts are powerful techniques, but they are ultimately simplifications of a publication’s full scientific contribution. A novel concept might be discussed in the main body of a paper but not mentioned in the abstract, and thus would be missed by our analysis. Furthermore, LDA identifies probabilistic themes, and the assignment of a single “dominant topic” to a paper is a necessary simplification for certain analyses; many papers are inherently multi-thematic.
- Citation Lag and Meaning: Citation counts are a widely accepted proxy for scientific impact but are subject to a significant time lag. Papers published late in the study period (e.g., 2024-2025) have not had sufficient time to accumulate citations, which may affect analyses comparing impact over time. Moreover, citations are not always a pure measure of positive impact; they can be perfunctory, negative, or self-serving, though on a large scale they are generally accepted as a valid indicator of visibility and influence.

#### Interpretation Boundaries

- Correlation vs. Causation: This study identifies numerous strong correlations, but the observational nature of bibliometric data means we cannot establish causation. For example, while we found a statistically significant correlation between open data sharing and higher citation counts, we cannot definitively conclude that sharing *causes* the increase in citations. It is also plausible that higher quality research is both more likely to be cited and more likely to be shared openly by confident authors. Similarly, the lead-lag relationships between Kaggle and academia show a temporal sequence but do not prove that one directly causes the other; both may be responding to a third, unobserved factor, such as a breakthrough in an underlying technology.

These limitations were mitigated where possible through the use of robust statistical methods, transparent reporting, and cautious, hedged language in our interpretations (e.g., “suggests,” “appears to correlate with”). We believe that by acknowledging these constraints, the findings of this study can be properly contextualized, providing a valuable and reliable, though not infallible, map of the medical genomics data science field.

## 5. Conclusion and Future Directions

This study set out to construct a comprehensive, data-driven map of the medical genomics data science field, aiming to illuminate its intellectual structure, social dynamics, and evolutionary trajectory. To achieve this, we employed a novel bibliometric framework that integrated evidence from two distinct ecosystems: the formal academic literature indexed in the Dimensions database and the practical, problem-solving environment of the Kaggle platform. This dual-lens approach enabled a unique synthesis of the theoretical and applied frontiers of the discipline.

The analysis revealed a field defined by rapid, maturing growth, organized around a robust, small-world network of collaborative research groups. We charted the conceptual evolution of the discipline from a collection of broad intersecting ideas to a dense, integrated network centered on the paradigm of deep learning. Our findings demonstrated that while scientific impact is democratically distributed across all major research themes, it is significantly amplified by a commitment to open data sharing. Crucially, by bridging academia and practice, we identified the role of competitive data science platforms as early indicators and potential accelerators for cutting-edge techniques, while also uncovering clear translational gaps that represent opportunities for future synergy.

These findings provide the first, to our knowledge, integrative map of both the theoretical and practical dimensions of medical genomics data science. By moving beyond a purely academic view, this work offers a more holistic understanding of the entire knowledge lifecycle; from data generation and foundational research to the application of state-of-the-art methods on practical challenges. This comprehensive perspective is vital for navigating a field that is becoming increasingly central to the future of precision medicine, providing a structured evidence base for researchers, institutions, and funding bodies to make more informed strategic decisions. The insights gleaned from this study offer several actionable takeaways: researchers can better position their work within the field’s thematic landscape, institutions can identify strategic areas for investment, and educators can align curricula with the demonstrated evolution of in-demand techniques.

Based on the observed trends, the field of medical genomics data science is poised to continue its trajectory of rapid innovation and increasing specialization. We anticipate a deeper integration of multi-modal data and the continued absorption of cutting-edge architectures from the broader machine learning community. As the discipline matures, fostering the feedback loop between practical challenges and academic inquiry will be paramount to accelerating the translation of genomic insights into clinical impact.

Looking forward, this study opens several avenues for future research. A longitudinal follow-up is necessary to track the evolution of the “bursting” keywords identified here and to validate the long-term forecasts of our growth models. Methodologically, future work could extend this analysis by incorporating patent data and clinical trial registrations to map the full “bench-to-bedside” pipeline, tracing the journey of a concept from publication to intellectual property and, finally, to clinical application. Comparative studies analyzing other data-intensive scientific fields; such as climate science or astrophysics; through a similar dual-lens framework could reveal universal patterns of knowledge diffusion in the age of open data and competitive science. Ultimately, by continuing to map the contours of this dynamic field, we can better navigate its complexities and harness its immense potential to redefine the future of medicine.

## Acknowledgements

The authors would like to thank the developers and maintainers of the open-access databases and tools used in this study, including Dimensions and Kaggle, for providing valuable resources that made this analysis possible. The author also acknowledges the guidance and support from peers and colleagues who provided feedback during manuscript preparation. No specific funding was received for this work.

## Notes

### Competing Interest Statement

The authors have declared no competing interest.

