## Supplementary Table S1 for "Bibliometric Insights into Medical Data Science Applications in Genomics: Evidence from Kaggle and Dimensions Datasets"

### S1 Table: Mapping Scheme for Kaggle Subfields to ANZSRC Research Fields

| Kaggle Subfield | Mapped ANZSRC Field | Rationale |
|---|---|---|
| Cancer Genomics | Clinical Sciences | Competitions in this subfield primarily focus on patient-level prediction tasks, such as tumor classification, metastasis detection, and treatment response, which directly aligns with the applied nature of Clinical Sciences. |
| Medical Imaging | Clinical Sciences | This subfield involves the analysis of clinical images (e.g., histopathology slides) for diagnostic purposes, placing it squarely within the domain of Clinical Sciences. |
| Variant Classification | Genetics | The core task is to determine the function and pathogenicity of genetic variations. This is a fundamental activity within the field of Genetics. |
| Epigenomics | Genetics | Epigenomics, the study of epigenetic modifications, is a sub-discipline of Genetics. The problems focus on predicting gene states based on modifications to DNA and histones. |
| Single-Cell Genomics | Cellular and Molecular Biology | Although clinically relevant, the data and analytical challenges (e.g., cell-type annotation, trajectory inference) are at the level of the individual cell, aligning with the focus of Cellular and Molecular Biology. |
| Proteomics | Biochemistry and Cell Biology | This subfield centers on the large-scale study of proteins, their functions, and structures, which is a core component of Biochemistry and Cell Biology. |
| Multi-Omics | Biochemistry and Cell Biology | The integration of different molecular layers (genomics, transcriptomics, proteomics) to understand biological systems is a key methodological approach in modern Biochemistry and Cell Biology. |
| Drug Discovery | Pharmacology and Pharmaceutical Sciences | Competitions in this area focus on predicting molecular activity, identifying inhibitors, and understanding mechanisms of action, all of which are central to Pharmacology and Pharmaceutical Sciences. |
| Pharmacogenomics | Pharmacology and Pharmaceutical Sciences | This subfield directly investigates how genetic variation influences drug response, making it a perfect fit for the Pharmacology and Pharmaceutical Sciences classification. |
